# A perturbation atlas of mouse neural organoids resolves macroglial reactivity and nominates drivers of degeneration

**DOI:** 10.64898/2026.10.09.757894

**Authors:** Jean-Samuel Dupré, Marina Signer, Jyoti Rao, Lukas Adam, Laura Gaspa-Toneu, Marina Almato-Bellavista, Irineja Cubela, Lorenca Sadiraj, Tobias Fuhrer, Julian Scherer, Teo Parés i Borrell, Manuel Schröter, Andreas Hierlemann, Mattia Zampieri, Matthias P. Lutolf, J. Gray Camp

**Affiliations:** Institute of Human Biology (IHB), Roche Pharma Research and Early Development, Roche Innovation Center Basel, Basel, Switzerland; Department of Biosystems Science and Engineering, ETH Zürich, Basel, Switzerland; Biozentrum, University of Basel, Basel, Switzerland; Department of Biomedicine, University of Basel, Basel, Switzerland; Institute of Bioengineering, School of Life Sciences and School of Engineering, Ecole Polytechnique Fédérale de Lausanne (EPFL), Lausanne, Switzerland

## Abstract

Understanding neuro-glial roles in homeostasis and disease requires scalable central nervous system (CNS) models, particularly as macroglia like astrocytes and Müller glia remain challenging to study in slow-developing human systems. Here, we generate and map the cellular diversity of mouse neural organoids spanning multiple brain regions and the retina. Benchmarking transcriptomic maturation against *in vivo* references captures postnatal neuro-macroglial states within a month. We show that macroglia are highly responsive to inflammatory cues and adopt specific transcriptomic states also found *in vivo*. Through a single-cell perturbation screen targeting 13 Müller-glia-specific pathways, we map how certain inflammatory cytokines drive degeneration-like stress programs in photoreceptors and distinct reactive states in Müller glia, including an Oncostatin M-driven chaperone-associated state. Mechanistically, we identify gp130-STAT3 signaling as a targetable driver of macroglial reactivity and photoreceptor distress. Together, these single-cell atlases provide a foundation for dissecting cellular crosstalk and neuro-glial pathology in the mature CNS.

## Main

The architecture of the mammalian central nervous system (CNS) is established through conserved, precisely timed morphogenetic processes ^1,2^. Across both the brain and the retina, progenitor cells migrate and differentiate to establish laminated structures, with macroglia (e.g., astrocytes and Müller glia) emerging among the latest cell types ^3,4^. Once established, the continuous crosstalk between neurons and macroglia is essential for sustaining the homeostatic environment required for sensory processing and cognitive functions. Breakdown of this cellular equilibrium contributes to CNS pathologies, with disease-associated glial states participating in disease progression by mediating synaptic dysfunction, metabolic breakdown, and eventually neuronal death ^5–8^. Features of brain neurodegeneration often manifest in the retina ^9^, and both organs share pathological hallmarks ^10,11^; yet, the mechanisms governing neuro-glial pathology remain poorly understood. Dissecting disease-associated macroglial states and advancing other frontiers in neurobiology require experimental systems capable of modeling mature mammalian neural tissue at scale.

Neural organoids are three-dimensional *in vitro* tissues that can be guided towards different CNS regions using molecular cues. They complement animal models because of their amenability to chemical and genetic manipulation, highcontent phenotyping, and high-throughput screening ^12–14^. However, the majority of current neural organoid studies are performed using human pluripotent-stem-cell-derived organoids and are therefore limited by the slow developmental clock inherent to human cells. Indeed, while these models have been successfully used to study early developmental processes such as neuronal patterning ^14–16^, cell-lineage decisions ^17–19^, and neurodevelopmental disorders ^20–23^, they struggle to efficiently reach late-stage, postnatal states without low-throughput interventions such as *in vivo* transplantation ^24–27^, or maintaining cultures for prolonged periods spanning years ^28^.

Given that developmental pacing varies significantly across species ^1,2,29^, non-human mammalian organoids offer a strategy to circumvent the prolonged maturation periods of human tissues. For instance, by leveraging the accelerated murine developmental clock, neural organoids derived from mouse stem cells expand the current *in vitro* toolkit, finding applications in protocol optimization ^30–33^, evolutionary studies ^34^, cross-species comparisons ^35,36^, and neuronal circuit assembly ^37,38^. They also open new avenues in iterative bioengineering, *in vitro*-to-*in vivo* benchmarking against genetically matched mouse models, co-culture with primary immune cells, and arrayed perturbation screens; altogether advancing 3R (Replacement, Reduction, Refinement) efforts by providing a scalable *in vitro* alternative to animal models.

Here, we establish an integrated atlas of mouse neural organoids spanning diverse regional identities to explore mature mammalian CNS physiology. Through dense multimodal time-course profiling, we demonstrate that mouse neural organoids recapitulate the accelerated pace of murine *in vivo* development, organizing into three-dimensional tissues containing mature macroglia and neurons in less than five weeks. Leveraging mouse retinal organoids, we execute a multiplexed, single-cell perturbation screen to map the macroglial phenotypic space. This atlas reveals distinct, cytokine-specific reactive macroglial states and a cytokine-induced degeneration-like state in photoreceptors. Finally, we show that the emergence of these states is driven by the gp130–STAT3 signaling axis.

## Results

### Mouse neural organoids generate diverse, matured neuronal and macroglial subtypes across CNS regions

We established a suite of protocols to generate neural organoids from mouse stem cells (Fig. 1a, Extended Data Fig. 1a). Organoids spanning cortical, thalamic, mesencephalic, cerebellar, retinal and mixed regional identities were cultured by adapting existing human and mouse neural organoid protocols ^20,39–44^, adjusting the timelines to reflect murine developmental rates when required ^2,29^. To enhance scalability, we tailored select protocols for automated, 96-well culture (Extended Data Fig. 1b-d, Methods).

**Figure 1.**
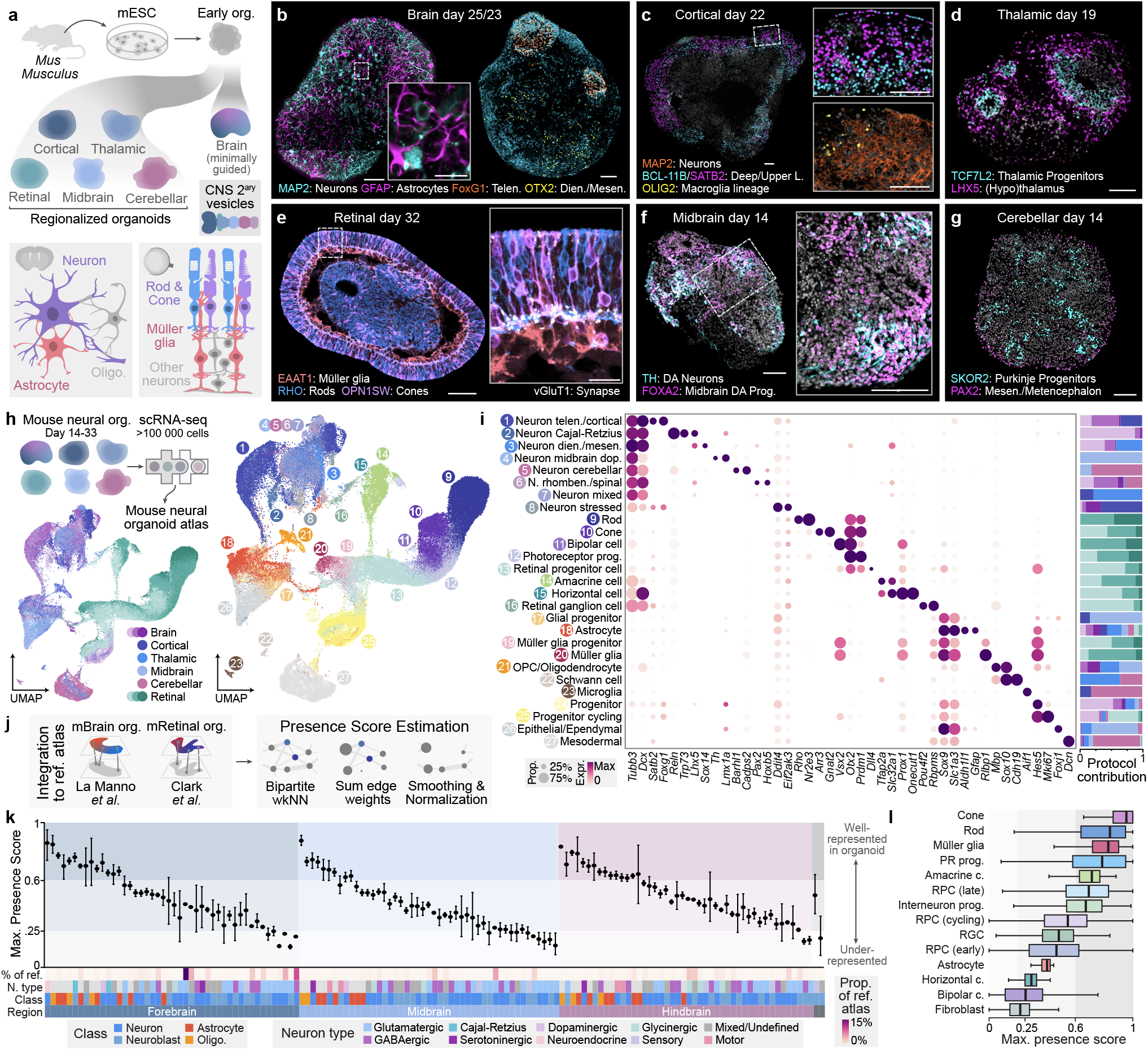
Mouse neural organoids generate diverse, matured neuronal and macroglial subtypes across CNS regions. **(a)** Conceptual workflow: organoids generated from mouse embryonic stem cells (mESCs) can be minimally guided towards brain identities or guided toward specific central nervous system (CNS) regions to investigate neuron-macroglia biology. **(b-g)** Representative immunostainings of neuronal, macroglial, and regional markers across regionalized mouse neural organoids. **(b)** Day 25 (left) and 23 (right) brain organoids with astrocytes (GFAP) interlaced between telencephalic (FoxG1) and diencephalic/mesencephalic (OTX2) neurons (MAP2). Scale bar inset: 25 *µ*m. **(c)** Cortical organoids containing deep (BCL-11B) and upper layer (SATB2) neurons (MAP2) alongside glial cells (OLIG2). **(d)** Thalamic organoids displaying thalamic progenitors (TCF7L2) and thalamic/hypothalamic neurons (LHX5). **(e)** Retinal organoids forming a stratified epithelium with rods (RHO), cones (OPN1SW), Müller Glia (EAAT1), and glutamatergic synapses (VGlut1) localized at the interface between cellular layers. Scale bar inset: 20 *µ*m. **(f)** Midbrain organoids displaying dopaminergic (DA) neurons (TH) and midbrain dopaminergic progenitors (FOXA2). **(g)** Cerebellar organoids featuring Purkinje neurons (SKOR2). Scale bars b-g: 100 *µ*m. **(h)** Top left: Single-cell RNA-sequencing (scRNA-seq) experimental overview. Bottom and right: UMAP embeddings of the integrated mouse neural organoid atlas colored by protocol (bottom), where increasing opacity reflects later time points, and cell type (right). **(i)** Dot plot of cell-type-specific marker expression (left) and stacked bar charts displaying protocols of origin for each cell type (right). **(j)** Computational strategy for benchmarking organoid fidelity against *in vivo* reference atlases: following the integration of brain and retinal organoid datasets with their respective reference atlases, a presence score is estimated for each reference cell type from the weighted sum of reference-organoid edges within a bipartite weighted k-nearest neighbor (wkNN) graph. **(k, l)** Maximum Presence Scores of primary tissue cell types in neural organoid datasets. **(k)** Brain organoids evaluated against La Manno *et al*. ^49^, highlighting recapitulation of diverse fore-, mid-, and hindbrain subtypes, together with macroglial (astrocyte and oligodendrocyte) clusters. **(l)** Retinal organoids evaluated against Clark *et al*. ^50^ exhibit strong fidelity for photoreceptor, Müller glia and interneuron subtypes. OPC = Oligodendrocyte progenitor cell; Rhomben. = Rhombencephalic; Prog. = progenitor; DA = dopaminergic; RPC = Retinal progenitor cell; RGC = Retinal ganglion cell; N. type = neuron type; PR = Photoreceptor.

Mouse neural organoids generated relevant, matured cell types including neurons and macroglia in less than 5 weeks of culture (Fig. 1b-g, Extended Data Fig. 1e-k). Minimally guided, multi-region brain organoids (hereafter referred to as brain organoids) displayed neuroglial interlacing, with astrocyte extensions interwoven around neuronal bodies and perisynaptic processes at neuronal interfaces, while oligodendrocytes wrapped neuronal axons (Fig. 1b, Extended Data Fig. 1l,m). Retinal organoids formed a laminated epithelium, where Müller glia spanned the tissue, interfacing with apically localized photoreceptors and basally localized interneurons (Fig. 1e, Extended Data Fig. 1k). Although retinal ganglion cells (RGCs) were observed early in development (Extended Data Fig. 1k), they did not persist at later time points, consistent with previous reports ^45–47^. Finally, we identified perineuronal nets, specialized extracellular matrix structures marking late-stage neural maturation ^48^, ensheathing neurons in older brain organoids from day 60 onwards (Extended Data Fig. 1n).

To construct a unified transcriptomic atlas resolving mouse neural organoid cellular composition, we performed single-cell RNA-sequencing (scRNA-seq) across time points and protocols (Fig. 1h). Integration and annotation revealed a diversity of matured cell types, including macroglia (astrocytes, oligodendrocytes, and Müller glia) and diverse excitatory and inhibitory neuronal subtypes whose regional identity aligned with the anatomical regions targeted by each respective protocol (Fig. 1h-i, Extended Data Fig. 2a-f). While minimally guided brain organoids exhibited broad regional diversity comprising predominantly forebrain and hindbrain neurons, guided protocols enriched for targeted identities, occasionally yielding cells from developmentally adjacent regions (e.g., *Foxg1*^+^ telencephalic and *En1*^+^ mesencephalic neurons in thalamic organoids). Beyond regional specification, cells expressed gene programs consistent with ongoing maturation. For instance, neurons broadly expressed the machinery required for active circuitry (*Slc12a5, Scn8a*) and visual cycle (*Sag, Abca4*), while macroglia acquired specialized support (*Slc1a3, Gja1, Rlbp1*) and myelinating profiles (*Mbp, Plp1*) (Extended Data Fig. 2g).

To assess the capacity of mouse neural organoids to recapitulate *in vivo* neuronal and macroglial cell type diversity, we integrated the generated single-cell datasets with reference transcriptomic atlases of the developing mouse brain ^49^ and retina ^50^ spanning E11-E18 and E11-P14, respectively. We computed presence scores ^25^ to quantify how well each primary tissue neuronal and macroglial type is captured in the organoid datasets (Fig. 1j, Supplementary Table 1). Brain organoids successfully recapitulated diverse fore-, mid-, and hindbrain identities, alongside macroglial (astrocytic and oligodendrocytic) subtypes, though certain prominent neuronal populations, notably forebrain GABAergic and mid-/hindbrain glutamatergic neurons, remained under-represented despite being abundant in the *in vivo* atlas (Fig. 1k). This mirrors limitations reported in human organoids, where certain brain regions and cell types, such as midbrain m1/m2-derived glutamatergic neurons, remain difficult to recapitulate ^25^. In parallel, retinal organoids showed the highest presence scores for rods, cones, Müller glia, and a subset of interneuron identities (amacrine cells) (Fig. 1l). Other cell types—such as early and cycling retinal progenitor cells (RPCs), astrocytes, retinal ganglion, horizontal, and bipolar cells—remained under-represented *in vitro*. This aligns with the advanced time points used for sequencing and reported developmental constraints across mammalian retinal organoids ^45^, coupled with mouse-specific limitations in *in vitro* bipolar cell generation ^51^.

To assess neuro-macroglial interactions, we performed Receptor-Ligand (R-L) analysis, revealing that expression of R-L pairs in organoids is consistent with primary *in vivo* tissue (cosine similarity ranging 0.72-0.92) (Extended Data Fig. 3a). Specifically, brain and retinal organoids recapitulated the compartmentalized expression of classic neuroglial pairs, including the glutamate transporters *Slc1a3* (EAAT1) and *Slc1a2* (EAAT2) expressed exclusively in macroglia (Extended Data Fig. 3b). Because macroglia handle 80-90% of CNS glutamate clearance ^52^, blocking this machinery with the broad-spectrum EAAT inhibitor L-trans-2,4-PDC (PDC) ^53^ should induce glutamate buildup and neuronal excitotoxicity ^54^. Indeed, PDC significantly reduced the area positive for neuronal-specific markers (MAP2 in brain organoids, RHO and OPN1SW in retinal organoids) while not significantly altering SOX9^+^ macroglial area (Extended Data Fig. 3c,d). Together, these data suggest that macroglia interact with neurons in organoids, notably to maintain neurotransmitter homeostasis.

### Multimodal trajectory mapping resolves accelerated maturation of mouse neural organoids

A defining feature of murine CNS development is its rapid pace relative to the human neurodevelopmental timeline ^2,29^. To determine how well mouse neural organoids recapitulate this accelerated timeline, we focused on brain and retinal organoids, executing a multi-modal temporal profiling encompassing protein immunostaining, scRNA-seq, metabolomics, and electrophysiology (Fig. 2a, Supplementary Table 2). In mouse brain organoids, time-course immunostaining resolved the rapid depletion of stem and neural progenitor pools (SOX2^+^) within the first two weeks, driven by a wave of neurogenesis (NeuN^+^, MAP2^+^) that peaked between days 8 and 17 (Fig. 2b,c). The emergence of macroglia after day 23 was evidenced by an increase in both astrocyte (10-FTHFDH^+^) and oligodendrocyte (SOX10^+^) markers. Time-course scRNA-seq profiling corroborated these global compositional shifts, capturing the reduced proportion of neural progenitors (21% of sequenced population) by day 14 and the progressive emergence of macroglial populations, comprising 16% of sequenced cells by day 30 (Fig. 2f). Aligning these changes in cell type composition to *in vivo* references ^49^ demonstrated that this day 14-30 *in vitro* window mirrors the compositional shifts occurring *in vivo* between embryonic days E8 and E18 (Fig. 2f). Of note, mouse brain organoids achieved this mature, macroglia-rich composition within 5 weeks, when their human counterparts require up to 20 weeks to reach equivalent stages (Fig. 2f). Similarly, time-course immunostaining in retinal organoids captured the rapid emergence of late-stage retinal cell types, including the expansion from day 21 onwards of rods (RHO^+^), cones (OPN1SW^+^), and Müller glia (EAAT1^+^) (Fig. 2d,e). Single-cell transcriptomics further confirmed the exhaustion of retinal progenitor pools, with retinal progenitor cells (RPCs) dropping from 55% of the population at day 21 to just 7% by day 33, concurrent with a massive expansion of rods, increasing from 1% of the sequenced cells on day 21 to 63% on day 33 (Fig. 2h). Of note, retinal organoids recapitulate the sequential emergence of cones before rods ^55^ (8% vs. 1% of sequenced cells on day 21) (Fig. 2d, 2h). Composition similarity analysis highlights that this day 21-33 *in vitro* window mirrors the shifts in cell type composition occurring *in vivo* around postnatal days P2-P8. In contrast, human retinal organoids require over 150 days *in vitro* to reach comparable compositional states (Fig. 2h).

**Figure 2.**
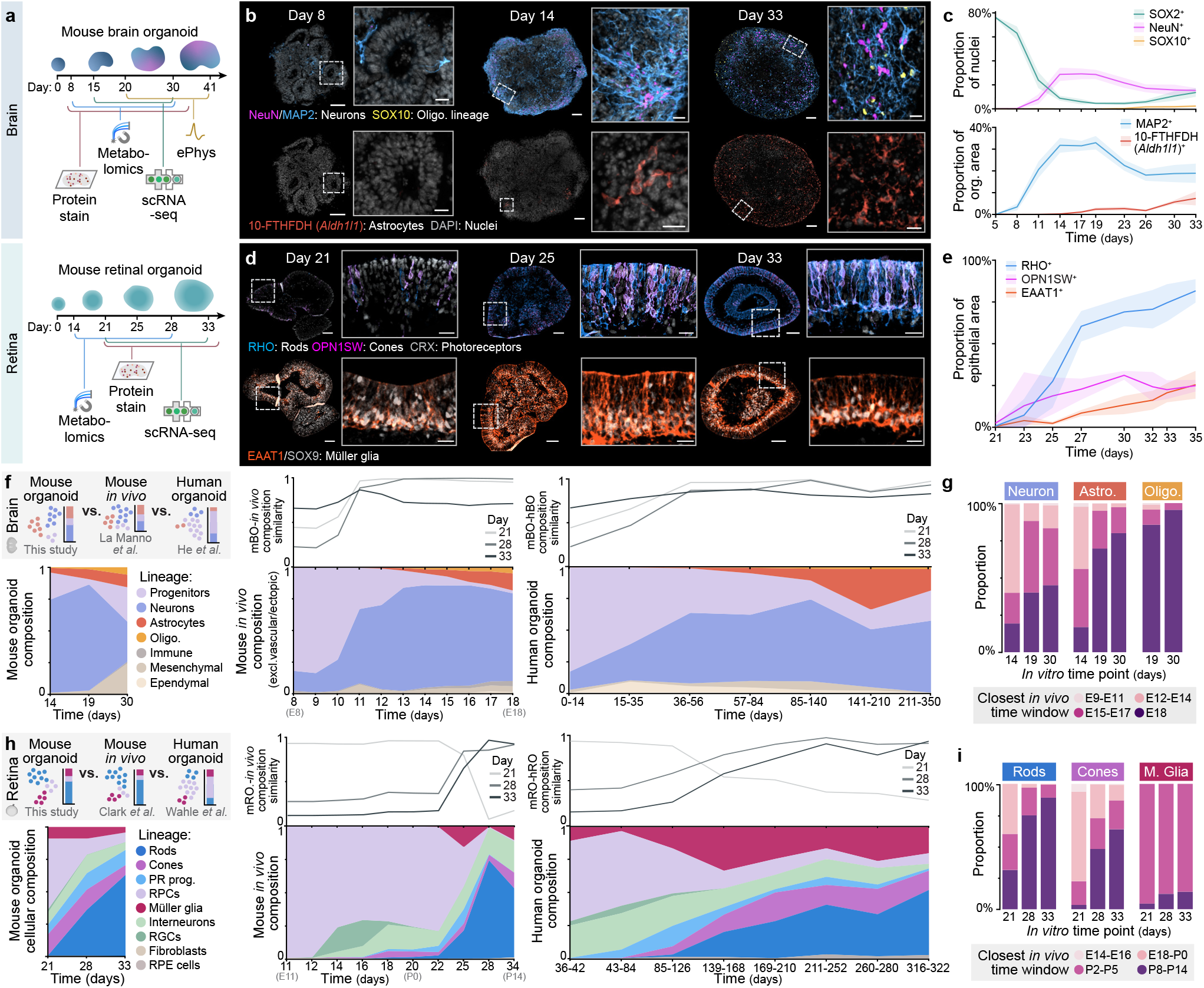
Multimodal trajectory mapping resolves accelerated maturation of mouse neural organoids. **(a)** Experimental workflow for the multi-modal time-course characterization of mouse brain and retinal organoids, integrating protein immunostaining, single-cell RNA-sequencing (scRNA-seq), whole-organoid metabolomics, and electrophysiology (ePhys). **(b)** Representative immunostainings of brain organoids at days 8, 14, and 33, highlighting the expansion of neurons (NeuN, MAP2) alongside the progressive emergence of oligodendrocytes (SOX10) and astrocytes (10-FTHFDH/Aldh1l1). Scale bars: 50 *µ*m; insets, 20 *µ*m. **(c)** Proportional abundance of cell type markers across brain organoid development. Representative images are displayed in Extended Data Fig. 5a. **(d)** Representative immunostainings of retinal organoids at days 21, 25, and 33, showing the development of rods (RHO), cones (OPN1SW), overall photoreceptors (CRX), and Müller glia (EAAT1, SOX9). Scale bars: 50 *µ*m; insets, 20 *µ*m. **(e)** Proportion of mouse retinal organoid epithelial area expressing photoreceptor and Müller glia markers over time. **(f-i)** Compositional and transcriptomic similarity trajectory mapping to *in vivo* and *in vitro* references. **(f)** Top left: Conceptual overview of cell type composition benchmarking against references. Bottom: Stream plots of cell type composition over time for mouse brain organoids (left), *in vivo* mouse brain (middle) and human guided or minimally guided brain organoids (right). Top middle and right: Similarity in composition between mouse organoids and respective reference. **(g)** Distribution of closest *in vivo* time points for mouse brain organoid neurons, astrocytes (astro.), and oligodendrocytes. **(h)** Top left: Conceptual overview of cell type composition benchmarking against references. Bottom: Stream plots of cell type composition over time for mouse retinal organoids (left), *in vivo* mouse retina (middle) and human retinal organoids (right). Top middle and right: Similarity in composition between mouse organoids and respective reference. **(i)** Distribution of closest *in vivo* time points for mouse retinal organoid rods, cones, and Müller glia (M. glia). RPE = Retinal Pigmented Epithelium.

We next projected organoid single-cell transcriptomes onto *in vivo* reference atlases ^49,50^ to predict equivalent developmental age. Both tissues showed clear, progressive temporal maturation of major cell types (Fig. 2g,i). In brain organoids, day 14 neurons mapped predominantly to intermediate embryonic stages (E12-E14), shifting toward late-embryonic states by day 30 (Fig. 2g). Late-emerging astrocytes and oligodendrocytes clustered at the latest time point (E18) in the reference atlas, suggesting that the transcriptomic maturation progresses to postnatal states. Retinal organoids exhibited similar dynamics, with day 21 photoreceptors aligning predominantly with perinatal stages (E18-P5) and steadily transitioning into advanced postnatal identities (P8-P14) by day 33, while Müller glia mapped exclusively to the latest reference points (P2-P14) (Fig. 2i).

To augment the transcriptomic trajectory mapping, we performed a dense bulk metabolomics time-course on brain and retinal organoids (Extended Data Fig. 4a). Analysis revealed a temporal progression from an early abundance of biosynthetic building blocks (e.g., membrane-building lipid metabolites in the brain and protein synthesis-related arginine-proline intermediates in the retina) to a subsequent increase in the relative abundance of neurotransmitters and their precursors by days 17-21 (Extended Data Fig. 4b-e,h-k). This increase in neurotransmitters parallels the emergence of neuronal populations (Fig. 2c,e) which aligns with the transcriptional upregulation of neurotransmitter genes (Extended Data Fig. 4f,l), and precedes the emergence of spontaneous electrophysiological activity in brain organoids (Extended Data Fig. 5d).

Crucially, late-stage organoids (days 25-30) exhibited specialized metabolic signatures indicative of glial maturation and support. In brain organoids, this was characterized by the progressive increase in the relative abundance of taurine, a macroglial-associated osmolyte and neuroprotectant ^56^, and lysophospholipids, bioactive lipids associated with oligodendrocyte specialization ^57,58^ (Extended Data Fig. 4e). Single-cell RNA-seq further confirmed the late-stage, macroglia-specific increase in expression of their corresponding biosynthetic enzymes at the transcriptomic level (Extended Data Fig. 4f). In contrast, late-stage retinal organoids exhibited increased levels of pentose phosphate pathway (PPP) intermediates (Extended Data Fig. 4k). While the PPP typically fuels nucleotide biosynthesis in dividing cells ^59^, its sustained activity in mostly post-mitotic retinal organoids suggests a pivot toward production of NADPH, a critical cofactor for both oxidative defense and the visual cycle ^60,61^. Supporting this hypothesis, scRNA-seq showed cell-type-specific upregulation of NADPH-dependent enzymes, including visual cycle components *Rdh8* and *Rdh12* in photoreceptors (Extended Data Fig. 4l,m). Together, these metabolic trajectories closely align with the transcriptomic mapping, reflecting a biochemical maturation that supports neuroglial functions.

To assess functional neuronal maturation, we tracked the spontaneous electrical activity of mouse brain organoids over the first six weeks of *in vitro* development. Consistent with the progressive transcriptional upregulation of voltage-gated ion channels and synaptic machinery (Extended Data Fig. 5b), high-density microelectrode array (HD-MEA) recordings revealed the rapid emergence of complex network activity in mouse brain organoids (Extended Data Fig. 5c). While most organoids were functionally silent on day 20, they developed spontaneous activity by day 27, which organized into synchronized network burst events by day 34. This was reflected by increases in overall firing rate, burst rate, and synchronicity over time (Extended Data Fig. 5d,e). Measured axonal conduction velocities spanned 0.1-1.5 m/s (Extended Data Fig. 5f), consistent with unmyelinated mammalian cortical axons ^62^. Acute application of the glutamatergic receptor antagonists AP5 and CNQX abolished this coordinated bursting and reduced the average firing rate by 53%, confirming that emergent network dynamics are driven by actively coupled, functional excitatory synapses (Extended Data Fig. 5g). While *in vivo* networks eventually desynchronize upon maturation of GABAergic interneurons and integration of sensory inputs ^63–65^, most organoids showed persistent, synchronous network activity through the recording period, likely reflecting the absence of these *in vivo* features ^37^. Crucially, mouse brain organoids achieve synchronized bursting within 4-5 weeks, whereas their human counterparts typically require 4 to 6 months of culture to reach equivalent milestones ^66–68^.

### Modulating CNS stress pathways reveals distinct macroglial responses across brain and retina

To dissect how macroglia and neurons respond to common neurodegeneration-linked stressors across the CNS, we performed single-cell transcriptomics of mouse brain and retinal organoids acutely exposed for 12 hours to interferons (IFN*α* + IFN*β* + IFN*γ*), inflammatory cytokines (TNF-*α* + IL-6 + IL-17A + IL-18 + OSM + SCF), and hydrogen peroxide (H_2_O_2_)-induced oxidative stress (Fig. 3a).

**Figure 3.**
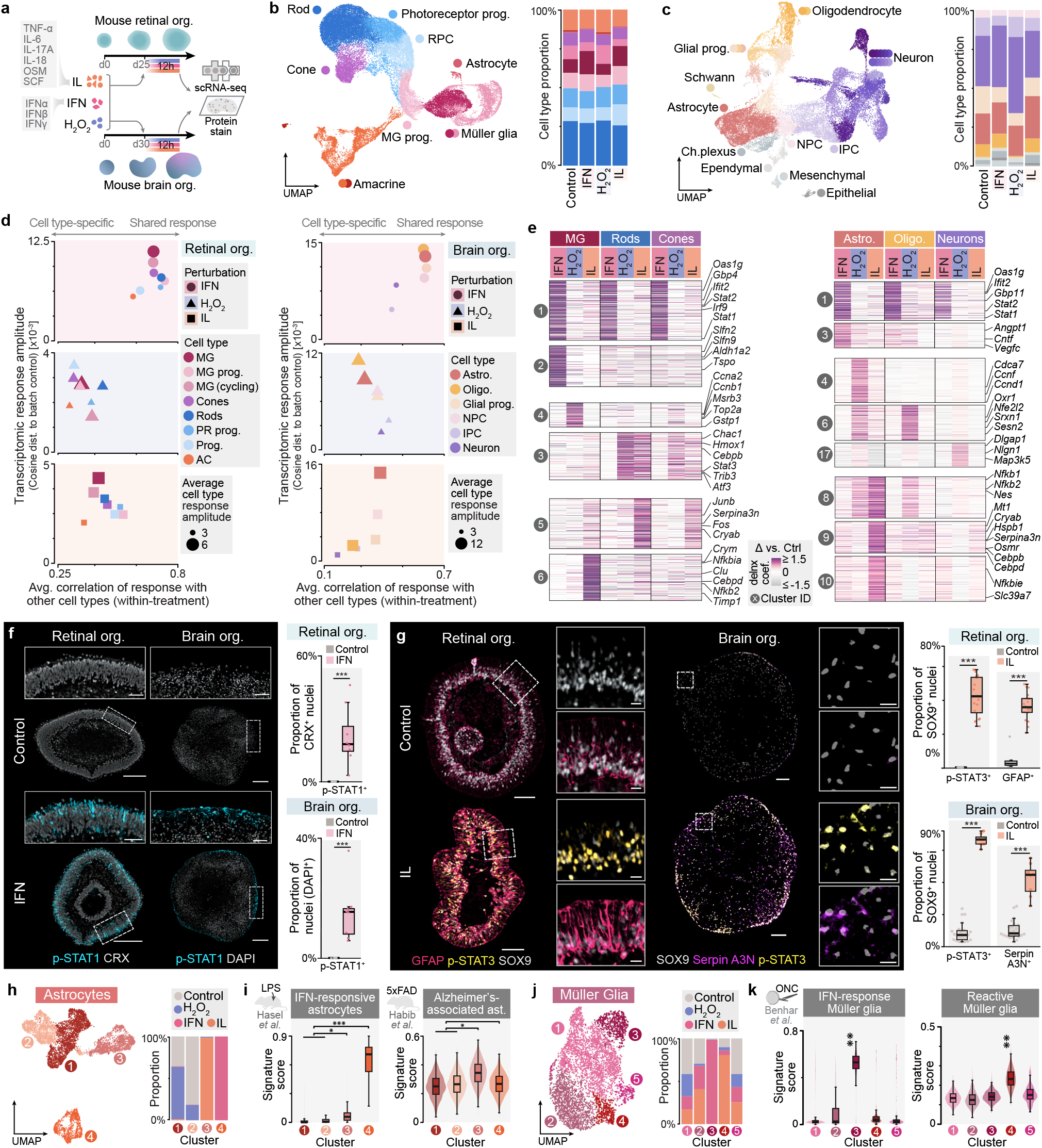
Modulating CNS stress pathways reveals distinct macroglial responses across brain and retina. **(a)** Experimental workflow for acute stress induction: mouse retinal and brain organoids were acutely exposed (12 h) to inflammatory cytokines (IL: TNF-*α*, IL-6, IL-17A, IL-18, OSM, SCF), interferons (IFN: IFN*α*, IFN*β*, IFN*γ*), or oxidative stress (H_2_ O_2_) prior to multi-modal profiling via single-cell RNA-sequencing and protein immunostaining. **(b, c)** UMAP embeddings (left) and cell type proportions across treatments (right) for retinal **(b)** and brain **(c)** organoids. **(d)** Scatter plots relating average transcriptomic response amplitude (cosine distance to batch control) with average within-treatment cross-cell type response correlation for retinal (left) and brain (right) organoids. **(e)** Heatmap displaying degree of change in expression (delnx effect size estimates) for differentially expressed genes across major cell types in retinal (left) and brain (right) organoids in response to IFN, H_2_ O_2_, and IL treatments, as compared to controls. Selected K-clusters and genes shown. **(f)** Representative immunostainings (left) and quantification (right) showing significantly increased proportion of phospho-STAT1^+^ nuclei in both retinal and brain organoids following IFN exposure. Points represent independent organoid replicates; *\*\*\** p < 0.001; Cohen’s h = 0.9 (retinal), 0.7 (brain). Scale bars: 200 *µ*m; insets, 50 *µ*m. **(g)** Representative immunostainings (left) and quantification (right) of IL-treated organoids showing significantly increased proportion of macroglial (SOX9^+^) nuclei positive for phospho-STAT3 and glial reactivity markers (GFAP in retinal Müller glia; Serpin A3N in brain astrocytes). Points represent independent organoid replicates; *\*\*\** p < 0.001; Cohen’s *h*_p-STAT3_ = 1.5 (retinal), 1.7 (brain); *h*_GFAP_ = 1.0; *h*_SerpinA3N_ = 0.9. Scale bars: 100 *µ*m; insets, 20 *µ*m. **(h-k)** High-resolution subclustering of macroglia and benchmarking against *in vivo* signatures. **(h, j)** UMAP embeddings (left) and treatment distribution (right) of astrocytes **(h)** and Müller glia **(j)** reveal distinct, perturbation-driven states. **(i, k)** Distribution of *in vivo* signature scores per macroglial subcluster for **(i)** astrocyte clusters 4 and 3 versus baseline clusters (1 and 2) for the ‘IFN-responsive’ and ‘Alzheimer-associated’ signature respectively (Cohen’s d = 7.2 and 1.1). **(k)** Müller glia clusters 3 and 4 versus baseline clusters (1, 2 and 5) for the ‘IFN-response’ and ‘reactive’ signature respectively (Cohen’s d = 2.7 and 2.7). Effect size: ✳✳✳ d > 3, ✳✳ d > 2, ✳ d > 1. MG = Müller glia; Prog. = progenitor; NPC = Neural progenitor cell; IPC = Intermediate progenitor cell; Ch. Plexus = Choroid plexus; PR = Photoreceptor; AC = Amacrine cells; LPS = lipopolysaccharide; ONC = Optic nerve crush.

Although all perturbed organoids contained the expected major cell types (Fig. 3b,c, Extended Data Fig. 6a-c), measuring response amplitudes and cross-cell type correlations uncovered specific patterns in the tissue stress response. Indeed, while all types mounted a correlated response to interferons (average Pearson’s r = 0.70 and 0.57) for retinal and brain organoids), the transcriptomic responses to inflammatory and oxidative stressors were more cell-type specific, with average correlations spanning 0.29-0.46 (Fig. 3d, Extended Data Fig. 6d). Furthermore, the distinct cell types responded with varying magnitudes; for instance, IL-exposed brain astrocytes exhibited a 15-fold larger cosine distance to batch control compared to their neuron counterparts (Fig. 3d). To dissect the programs driving these responses, we performed differential expression and gene set enrichment analyses (GSEA) across the major cell types. This revealed compartmentalized transcriptomic modules, notably isolating macroglial-specific reactive programs (e.g., retinal k-cluster 6 and brain k-cluster 9) from neuronal-specific ones (e.g., retinal k-cluster 3 and brain k-cluster 17) (Fig. 3e, Extended Data Fig. 6e,f).

Interferons, notably secreted by microglia ^69,70^, regulate response to infections and are implicated in CNS injury and neurodegeneration ^71,72^. Upon exposure to interferons, brain and retinal organoids mounted a correlated response across cell types (average Pearson’s r = 0.70 and 0.57 for retinal and brain organoids), with macroglia consistently displaying the strongest response (Fig. 3d). Cells across both tissues upregulated genes associated with interferon signaling and the JAK-STAT signaling pathway, including antiviral transcription factor components (*Stat1, Stat2, Irf9*) and classical interferon-stimulated genes (*Oas1g, Ifit2, Gbp4*) (Fig. 3e, Extended Data Fig. 6f). Confirming STAT1 complex activation, IFN exposure drove a significant increase in phospho-STAT1^+^ nuclei across both tissues (p < 0.001) (Fig. 3f). In addition, both Müller glia and astrocytes upregulated multiple ligands involved in immune recruitment and activation (*Cxcl10, Csf1, Ccl2, H2-Aa*) (Extended Data Fig. 6g).

Oxidative stress is a driver of neurodegeneration which can disrupt neuro-glial communication ^73,74^. In contrast to interferons, transcriptomic shifts induced by hydrogen peroxide were more cell-type-specific (Pearson’s r = 0.35 for retinal and brain organoids) (Fig. 3d, Extended Data Fig. 6d,h, Supplementary Table 3). Retinal organoids mounted a comparatively muted global response lacking significantly enriched GSEA terms, whereas brain organoids exhibited strong transcriptomic shifts driven primarily by macroglia (Fig. 3d, Extended Data Fig. 6d,f). Retinal Müller glia upregulated oxidative stress protection genes (*Gstp1, Msrb3*) alongside cell division markers (*Top2a, Ccnb1, Ccna2*), consistent with a mild expansion of cycling cells (from 8.6% of sequenced cells in control to 11.9% in H_2_O_2_-treated) (Fig. 3b,e). Similarly, brain astrocytes and oligodendrocytes upregulated an NRF2-driven antioxidant program (*Srxn1, Sesn2, Oxr1*)—corroborated by increased levels of the NRF2 target HO-1 in astrocytes (Extended Data Fig. 6i)—alongside a suite of cell cycle regulators (*Ccnd1, Ccnf, Cdca7*), suggesting a possibly shared macroglial proliferative response to oxidative insult (Fig. 3e).

Latent inflammation, driven in part by macro- and microglia-secreted cytokines, is another hallmark of CNS disease ^11,75^. We selected TNF-*α*, IL-6, IL-17A, IL-18, OSM and SCF as relevant cytokines likely to model a broad inflammatory environment (Supplementary Table 4). Upon exposure to this combination of inflammatory cytokines (IL), both retinal and brain organoids elicited a robust, primarily macroglia-driven response characterized by global upregulation of inflammation-related pathways (Fig. 3d, Extended Data Fig. 6d,f). Both retinal Müller glia and brain astrocytes activated core JAK-STAT and NF-*κ*B networks, evidenced by the upregulation of downstream effectors (*Cebpd, Cebpb, Nfkb2, Nfkbie*) and significantly increased proportion of phospho-STAT3^+^ nuclei (p < 0.001) (Fig. 3f,g). Crucially, Müller glia and brain astrocytes shifted to a reactive glial state, a known phenotype observed *in vivo* upon injury or inflammation ^76–78^. Upregulated genes encompassed distinct modules governing cytoskeletal and matrix remodeling (*Gfap, Nes, Timp1*), immune response regulation (*Serpina3n, Osmr, Lcn2*), chemokine signaling (*Cxcl1/10, Ccl2/5*), metal ion homeostasis (*Mt1/2, Cp, Slc39a7*), neurotrophic support (*Fgf2, Ngf, Vgf, Bdnf*) and proteostatic chaperone activity (*Clu, Cryab, Hspb1*) (Fig. 3e, Extended Data Fig. 6g). Significant induction of GFAP in Müller glia and Serine protease inhibitor A3N (Serpin A3N) in astrocytes was validated at the protein level via immunostaining (p < 0.001) (Fig. 3g).

To determine if macroglial perturbation-driven identities recapitulate relevant *in vivo* states, we benchmarked them against gene signatures from *in vivo* mouse models of CNS injury or disease (Supplementary Table 5). Subclustering of astrocytes and Müller glia revealed distinct, treatment-specific states, including two IFN-specific clusters marked by *Stat1* and *Ifit1*, and two IL-specific clusters marked by *Serpina3n* and *Gfap* (Fig. 3h,j, Extended Data Fig. 6j). IFN-driven macroglial states have been described in mouse models of inflammation ^79^ and sustained injury ^80^, whereas reactive-like states are present in mouse models of degeneration ^6^ and acute injury ^80^. Comparing organoid macroglia to gene signatures from these models confirmed the similarity of treatment-induced states to these *in vivo* states. Indeed, the IFN-specific *in vitro* astrocyte cluster showed increased scores (Cohen’s d = 7.2)) for the signature derived from *in vivo* ‘interferon-responsive’ astrocytes ^79^, and the IL-driven *in vitro* state uniquely enriched d = 1.1 for the ‘disease-associated astrocyte’ signature derived from an *in vivo* Alzheimer’s model ^6^ (Fig. 3i). Likewise, the IFN-specific Müller glia *in vitro* cluster showed increased scores d = 2.7 against the signature derived from ‘IFN-response’ Müller glia *in vivo* ^80^, while the IL-driven *in vitro* state (Cluster 4) is specifically enriched d = 2.7 for the *in vivo* ‘Reactive Müller glia’ signature ^80^ (Fig. 3k). Altogether, these data demonstrate the capacity of mouse neural organoid macroglia to adopt distinct, *in vivo*-relevant states, which provides a foundation to investigate their emergence and functional consequences.

### Single-cell perturbation screen maps the macroglia phenotypic space and nominates pathways relevant in photoreceptor degeneration

Multiple brain astrocyte states have been described and linked to CNS disease progression ^6,58,79,81–83^, however the diversity of states that Müller glia can adopt remains poorly defined. To address this gap and map the Müller glia phenotypic space, we designed a multiplexed single-cell perturbation screen targeting 13 Müller glia-specific, disease-relevant pathways using 29 modulators in triplicates (Fig. 4a, Extended Data Fig. 7a and Supplementary Table 6). For each molecule we performed dose-optimization, selecting the highest concentration displaying no visual evidence of cytotoxicity (Extended Data Fig. 7b).

**Figure 4.**
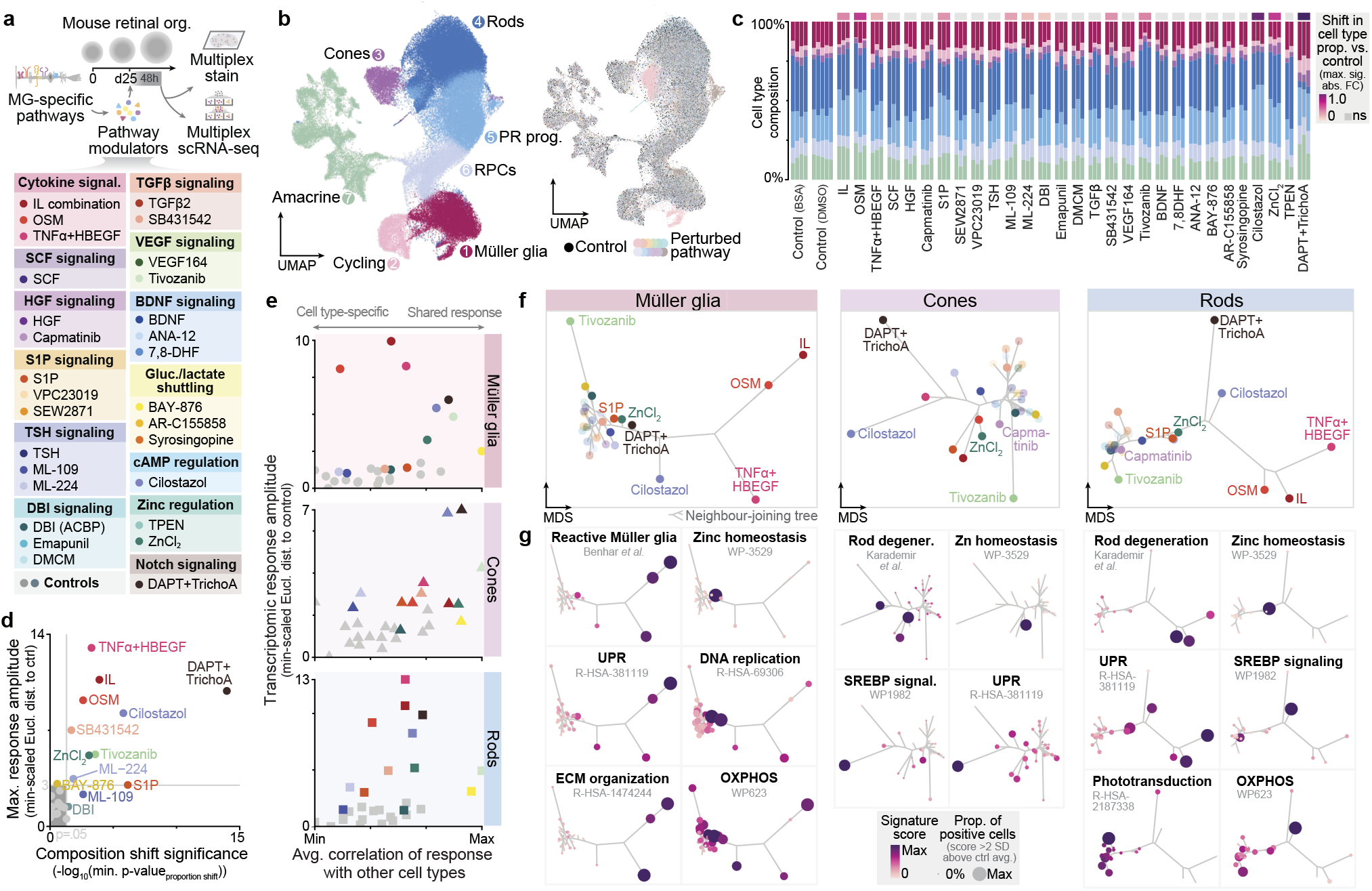
Single-cell perturbation screen maps the macroglia phenotypic space and nominates pathways relevant to neurodegeneration. **(a)** Experimental workflow for the multiplexed single-cell perturbation screen. Differentiated (day 25) mouse retinal organoids were exposed for 48 hours to a curated library of 29 small molecules and recombinant proteins targeting 13 distinct Müller glia-enriched signaling pathways, followed by multiplexed single-cell RNA-sequencing (scRNA-seq) and protein immunostaining. **(b)** UMAP embeddings of the integrated perturbation atlas colored by cell type (left) and targeted pathway (right). **(c)** Cell type composition across biological replicates for all perturbations and vehicle controls (BSA, DMSO). The colored tiles (top) indicate the maximum significant (p < 0.05) absolute fold-change (FC) in cell type proportions versus controls. **(d)** Scatter plot relating compositional shift significance (minimum log-transformed p-value from proportion shift testing) to the maximum transcriptomic response amplitude (min-scaled Euclidean distance to control) across perturbations. Only perturbations either inducing a significant shift in cell type composition (p < 0.05) or a large (>3) maximum response amplitude are highlighted. **(e)** Scatter plots relating transcriptomic response amplitude (min-scaled Euclidean distance to control) to the average correlation in transcriptomic response for major cell types, colored by treatment. Low-impact treatments as determined in (d) are greyed out. **(f)** Multidimensional scaling (MDS) embeddings with projected neighbor-joining (NJ) trees (phylomorphospaces) visualize the phenotypic distances between perturbation-induced cellular states across major cell types. **(g)** Phylomorphospaces from (f) overlaid with average signature scores identify specific transcriptomic modules associated with distinct regions of the phenotypic space. Dot color reflects average signature score from defined gene sets. Dot size denotes the zero-max-normalized fraction of positive cells (>2 SD above the control mean). Gluc. = Glucose; cAMP = cyclic AMP; TrichoA = Trichostatin A; Eucl. dist. = Euclidean distance; Degener. = Degeneration.

Following quality control and cell-type annotation of the resulting 115,000 cell perturbation atlas (Fig. 4b, Extended Data Fig. 7c), we assessed the impact of each perturbation on both organoid composition and transcriptomic state across three independent biological replicates per condition. Overall, twelve perturbations significantly altered cellular proportions, impacting principally rods, cones, Müller glia and cycling populations (Fig. 4c,d, Extended Data Fig. 7d). Notably, the PDE3 inhibitor Cilostazol, sphingosine-1-phosphate (S1P), inflammatory cytokines (IL) and Oncostatin M (OSM) led to a significantly decreased proportion of rods (p < 0.05) (Fig. 4c, Extended Data Fig. 7d).

To quantify the amplitude of the response across perturbations, we calculated the Euclidean distance between control and perturbed transcriptomes for each cell type (Extended Data Fig. 7e). Treatment with TNF-*α* and HBEGF (TNF*α*+HBEGF) ^84,85^, OSM, IL, a combination of the Notch inhibitor DAPT ^86^ with Trichostatin A ^87^, Cilostazol, the VEGFR inhibitor Tivozanib, zinc chloride (ZnCl_2_), and the Activin/TGF-*β* inhibitor SB431542 yielded the strongest responses in any cell type (Fig. 4d). Correlation in transcriptomic response also varied between treatments; for instance, Cilostazol induced a more correlated transcriptomic shift across cell types, while OSM provoked a large but weakly correlated response in Müller glia and rods (Pearson’s r=0.22 and 0.32 respectively) (Fig. 4e).

To visualize this complex phenotypic space, we constructed multidimensional scaling (MDS) plots with projected neighbor-joining trees for the main cell types based on differential gene expression, where distances reflect how distinct the responses between treatments are (Fig. 4f). In line with our previous analyses, most treatments which had shown a weak response clustered closely together, while the treatments with high response amplitude occupied distinct ‘pockets’ of the MDS space. Combining this topological approach with gene set enrichment analysis (GSEA) revealed several specific transcriptomic modules governing retinal response to perturbation (Fig. 4g, Extended Data Fig. 7f).

For instance, ZnCl_2_ exposure specifically triggered a metal-detoxification program across cell types, upregulating metallothioneins (*Mt1, Mt2*) and the zinc transporter *Slc30a2*, alongside activation of the Unfolded Protein Response (UPR; *Hspa5, Ddit3, Ppp1r15a*) (Fig. 4f,g, Extended Data Fig. 7f,g). The ability to capture this metal-handling signature demonstrates the sensitivity of the screen to resolve precise, pathway-specific response programs.

In parallel, DAPT+Trichostatin A shifted 77-93% of Müller glia into a reprogramming-like state characterized by upregulation of the transdifferentiation markers *Ascl1* and *Neurog2*, as well as synaptic genes such as *Arpp21* and *Ctnna2* ^88,89^ (Extended Data Fig. 8a-d), in line with reports previously showing that DAPT and Trichostatin A impact Müller glia transdifferentiation in mice ^86,87^. DAPT+Trichostatin A and S1P treatments induced DNA replication programs in Müller glia but not in photoreceptors, in line with the observed expansion of cycling cells (2.0- and 1.5-fold vs. control for DAPT+Trichostatin A and S1P) (Fig. 4f,g, Extended Data Fig. 7d,f,g).

The PDE3 inhibitor Cilostazol triggered upregulation of key genes of the Integrated Stress Response (ISR) amino acid branch ^90^ (*Asns, Sars, Aars*), and genes involved in oxidative phosphorylation (OXPHOS) across cell types (Fig. 4f, Extended Data Fig. 7f,g). This is consistent with effects of PDE inhibition in murine retinal degeneration models ^91,92^. On the other hand, c-Met inhibitor Capmatinib, together with Cilostazol, specifically upregulated sterol metabolism programs (notably SREBP targets *Hmgcr, Fads2, Cyp51, Scd1*) in rods and cones but not in Müller glia (Fig. 4f, Extended Data Fig. 7f,g).

Interestingly, Activin/TGF-*β* inhibition via SB431542 induced an increase in the proportion of cones (1.5-fold vs. control) (Extended Data Fig. 7d,e), and shifted photoreceptor progenitors to a new, specific state (Extended Data Fig. 8e-k). Fate transition probability analysis confirmed that SB431542treated progenitors exhibit a higher probability of transitioning into cones at the expense of rods (Extended Data Fig. 8i,j). Molecularly, these SB431542-specific progenitors repressed expected SMAD/TGF-*β* targets (*Smad3/4, Pmepa1*) alongside master rod determinants (*Nrl, Nr2e3*), while selectively upregulating cone specification markers (*Thrb, Rxrg, Arr3*) (Extended Data Fig. 8k).

Finally, three cytokine-mediated perturbations (IL, OSM, and TNF*α*+HBEGF) drove Müller glia into reactive states characterized by glial reactivity markers (*Gfap, Vim, Col12a1*) and increased ‘Reactive Müller glia’ signature scores ^80^ (Fig. 4f,g, Extended Data Fig. 7f,g). Furthermore, these cytokines induced upregulation of degeneration-associated gene programs in photoreceptors, leading to increased ‘Photoreceptor Degeneration’ signature scores ^93^, together with downregulation of phototransduction programs, particularly in rods (Fig. 4f,g Extended Data Fig. 7f,g). This dual phenotype implicates the downstream signaling networks, principally JAK/STAT, NF-*κ*B, PI3K/Akt, ERK and p38/JNK ^94–96^, as potential drivers of retinal degeneration. We therefore directed our focus toward deciphering the states and signaling pathways induced by OSM, IL and TNF*α*+HBEGF.

### Oncostatin M induces a distinct reactive state in Müller glia and a degeneration-like state in photoreceptors via the gp130-STAT3 signaling axis

Subclustering of Müller glia revealed 5 molecularly distinct treatment-induced states (Extended Data Fig. 8a-d). In particular, while both OSM and TNF*α*+HBEGF induced common changes in Müller glia, including the downregulation of homeostatic markers (*Rlbp1, Glul*), upregulation of intermediate filaments (*Gfap, Vim, Nes*) and STAT3 targets (*Cebpb, Stat3*), these treatments led to two distinct reactive states (Reactive I/II; Fig. 5a-c, Extended Data Fig. 8a-d). The combination of TNF-*α* and HBEGF, previously shown to induce glial reactivity and photoreceptor distress in organoid models ^84,85^, drove Müller glia into a proliferative, pro-inflammatory ‘Reactive I’ state. This state was uniquely defined by the upregulation of genes involved in neuroinflammatory signaling (*Lrrk2, Cxcl1*), cell cycle progression (*Ccnd2*), and extracellular niche modulation (*Kitl, Dkk2, Adamtsl3*) (Fig. 5b, Extended Data Fig. 8b). In contrast, OSM exposure drove Müller glia into an alternative ‘Reactive II’ state enriched for a distinct, chaperone-associated proteostatic program. Specifically, OSM uniquely upregulated core chaperones (*Clu, Cryab, Hspb1*), neurotrophic factors (*Ntf3, Ngf, Fgf2*), and acute-phase AP-1 complex members (*Fos, Jun, Atf3*), alongside other STAT3 targets (*Osmr, Socs3*) including inflammatory cytokines (*Ccl2, Cxcl5*) (Fig. 5b, Extended Data Fig. 8b). This divergent pathway activation was validated at the protein level where both cytokine perturbations triggered significant increases in STAT3 phosphorylation and GFAP accumulation (p < 0.01), while the induction of the chaperone HspB1 was specific to OSM exposure (p < 0.001) (Fig. 5c, Extended Data Fig. 9a). Of note, the inflammatory cytokine combination (IL) drove Müller glia into the ‘Reactive II’ state in similar proportions (68-77% of total) as OSM alone (59-84% of total) (Fig. 5b), and systematic testing of the IL components confirmed that OSM is the dominant driver of this specific glial reactivity phenotype (Extended Data Fig. 9b).

**Figure 5.**
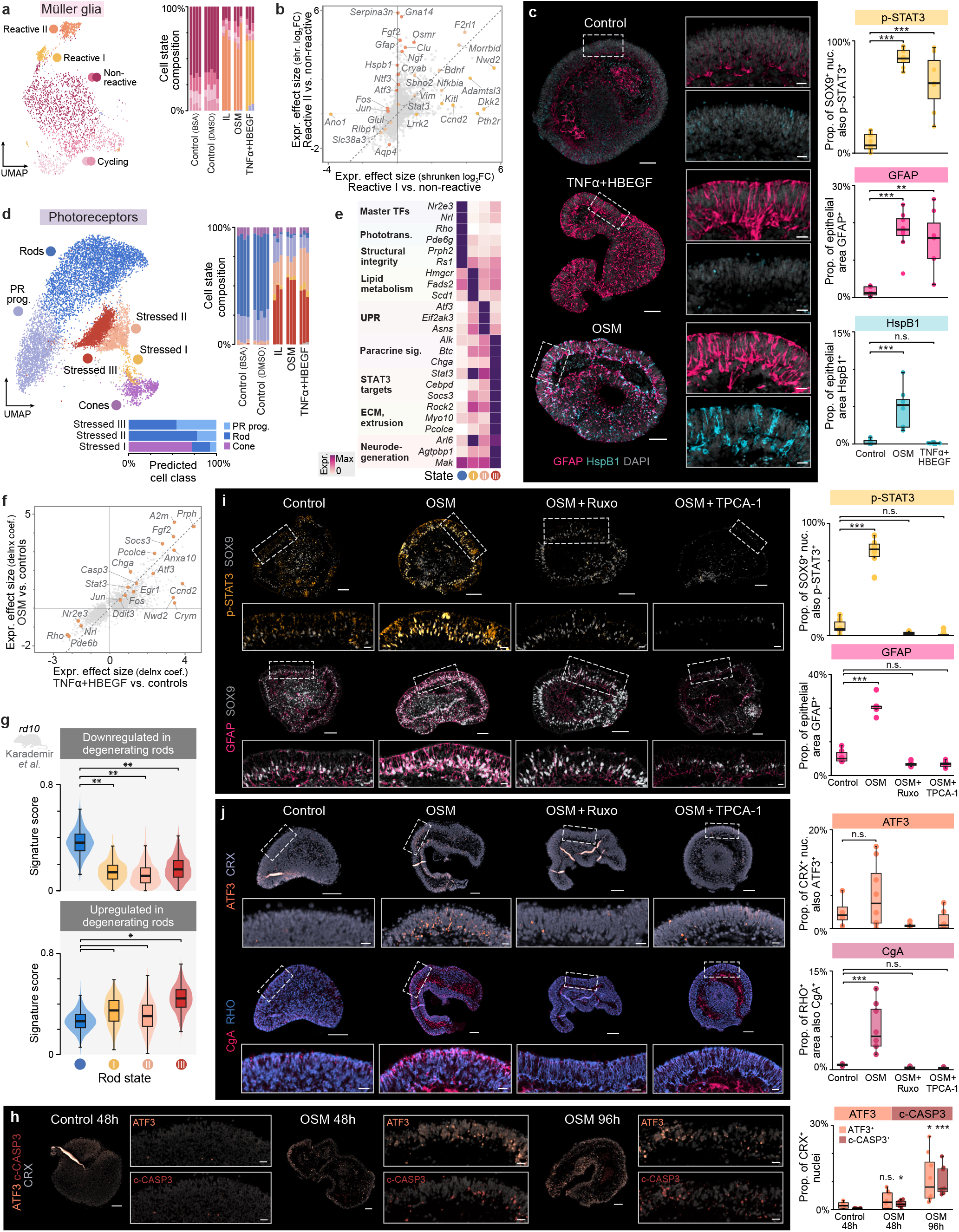
Oncostatin M induces a distinct reactive state in Müller glia and a degeneration-like state in photoreceptors via the gp130–STAT3 signaling axis. **(a)**UMAP embeddings (left) and distribution of cell states (right) for Müller glia from the IL, OSM, TNF*α*+HBEGF, and control conditions. 61–66% of TNF*α*+HBEGF-treated Müller glia were in the “Reactive I” state, while 59–84% of IL-treated and 68–77% of OSM-treated Müller glia were in the “Reactive II” state. **(b)** Scatter plot relating the degree of change in expression (shrunken log_2_ fold-changes) of differentially expressed genes in “Reactive I” and “Reactive II” states relative to non-reactive Müller glia. **(c)** Representative immunostainings (left) and quantification (right) of mouse retinal organoids treated with TNF*α*+HBEGF or OSM for 48 h, showing significantly increased glial levels of GFAP and phospho-STAT3 in both conditions, but increased HspB1 only in OSM-treated organoids. Points represent independent organoid replicates; *\*\*\** p < 0.001, *\** p < 0.05; Cohen’s *h*_p-STAT3_ = 1.31/1.80, *h*_GFAP_ = 0.54/0.63, and *h*_HspB1_ = −0.02/0.35 for OSM/TNF*α*+HBEGF. Scale bars: 100 *µ*m; insets, 20 *µ*m. **(d)** UMAP embedding (left) and distribution of cell states (right) for photoreceptors from the IL, OSM, TNF*α*+HBEGF, and control conditions. Bottom: predicted cell-class composition for the stressed photoreceptor states. **(e)** Average zero-max-normalized gene expression of selected gene modules across rods and stressed photoreceptor states. **(f)** Scatter plot relating the degree of change in expression (delnx effect-size estimates) of differentially expressed genes in OSM- and TNF*α*+HBEGF-treated rods (predicted class) relative to control. **(g)** Distribution of *in vivo* rod-degeneration signature scores across rods and stressed photoreceptor states. While all three stressed states showed reduced scores for a set of genes downregulated (down.) during *in vivo* rod degeneration (Cohen’s d = −2.3, −2.7, and −2.1 for states I, II, and III versus rods), only the “Stressed III” state showed a large increase (effect size >1) for a set of genes upregulated (up.) during rod degeneration (Cohen’s d = 0.9, 0.4, and 2.0 for Stressed I, II, and III versus rods). Effect size: ✳✳✳ d > 3, ✳✳ d > 2, ✳ d > 1. **(h)** Representative immunostainings (left) and quantification (right) of the ER-stress marker ATF3 and apoptosis marker cleaved Caspase-3 (c-CASP3) in mouse retinal organoids exposed to OSM for 48 h and 96 h. Points represent independent organoid replicates; significance labels reflect comparisons against the respective controls for each marker; *\*\*\** p < 0.001, *\** p < 0.05; Cohen’s *h*_ATF3_ = 0.08/0.41, *h*_c-CASP3_ = 0.13/0.50 for 48 h/96 h; h = 0.36 and p < 0.001 for c-CASP3 at 96 h versus 48 h. Scale bars: 100 *µ*m; insets, 20 *µ*m. **(i-j)** Representative immunostainings (left) and quantification (right) for **(i)** macroglial reactivity markers and **(j)** photoreceptor stress markers in mouse retinal organoids exposed to OSM (48h), or OSM co-treated with the STAT3 inhibitor Ruxolitinib (Ruxo) or TPCA-1. Points represent independent organoid replicates; significance for increases relative to control was determined using a one-sided Welch’s t-test on arcsine-transformed proportions; *\*\*\*p <* 0.001; Cohen’s *h*_p-STAT3_ = 1.60/−0.26/−0.40, *h*_GFAP_ = 0.55/−0.13/−0.13, *h*_ATF3_ = 0.11/−0.20/−0.15, and *h*_CgA_ = 0.32/−0.07/−0.08 for OSM/OSM+Ruxolitinib/OSM+TPCA-1. Scale bars: 100 *µ*m; insets, 20 *µ*m.Phototrans. = Phototransduction; Paracrine sig. = Paracrine signaling; CgA = Chromogranin A

Beyond glial reactivity, OSM, along with the IL combination and TNF*α*+HBEGF, shifted photoreceptors into three distinct photoreceptor states (‘Stressed I/II/III’; Fig. 5d,e, Extended Data Fig. 8e-h). Unlike the divergent reactive trajectories observed in Müller glia, both OSM and TNF*α*+HBEGF induced these stressed photoreceptor states in comparable proportions, and only a minor subset of genes (e.g., *Ccnd2, Nwd2, Crym*) were uniquely enriched in the TNF*α*+HBEGF treatment (Fig. 5d,f). All three states were characterized by downregulation of photoreceptor marker genes (*Rho, Nrl, Nr2e3, Sag, Pde6b*) (Fig. 5e, Extended Data Fig. 7f). The first ‘Stressed I’ state was the only state comprising cells from the cone class (72% of total) and specifically upregulated genes involved in lipid homeostasis classically controlled by SREBPs (*Cyp51, Hmgcr, Fads2, Scd1*) (Fig. 5d,e). The second ‘Stressed II’ state exhibited an Unfolded Protein Response (UPR) driven by the PERK-ATF4/3 axis (*Eif2ak3, Atf4/3, Ddit3, Asns*) (Fig. 5e,f, Extended Data Fig. 8f). Finally, the third, ‘Stressed III’ state also exhibited a mild PERKATF4/3-driven UPR signature, but was in addition uniquely defined by the activation of paracrine signaling (*Chga, Alk, Fgf2, Btc*) and the JAK/STAT inflammatory axis (*Stat3, Cebpd, Socs3*) (Fig. 5e,f, Extended Data Fig. 8f). This state also upregulated ECM remodeling genes (*Rock2, Myo10, Scube1, Pcolce, Wwtr1*) potentially implicated in stressed photoreceptor extrusion ^85^. When compared to *in vivo* data, all three stressed photoreceptor states showed reduced signature scores for a set of genes downregulated during rod degeneration ^93^ (Fig. 5g; Cohen’s d = −2.3, −2.7, and −2.1 for states I, II, and III). Shared downregulated genes included rod marker genes such as *Nrl, Nr2e3* and *Rho* (Fig. 5e-f). However, only the ‘Stressed III’ state showed a large increase (effect size >1) in signature score for genes upregulated during *in vivo* rod degeneration ^93^ which include markers of neuronal degeneration (*Agtpbp1, Mak, Arl6*) (Fig. 5e-g; Cohen’s d = 0.9, 0.4, 2.0 for Stressed I, II and III). These results suggest that the ‘Stressed III’ identity aligns with *in vivo* transcriptomic features of rod degeneration.

Tracking OSM-driven glial and photoreceptor protein markers over a 4-day exposure showed significantly increased proportions of p-STAT3^+^ Müller glia and GFAP^+^ area within 12 hours of OSM exposure (p < 0.05), progressively increasing further through 48-96 hours (Extended Data Fig. 9c, Fig. 5j). In photoreceptors, the the proportion of cells positive for the ER stress marker ATF3 remained low and non-significant at 12h, increasing to significant levels at 96h (Fig. 5h, Extended Data Fig. 9c). The proportion of photoreceptors positive for the apoptosis marker cleaved Caspase-3 was significantly higher than in control organoids after 48h (p < 0.05) and further increased by 96h (p < 0.001) (Fig. 5h). Finally, the fraction of rod area (RHO^+^) positive for the intra-vesicular matrix protein Chromogranin A (CgA) was also significantly increased after 48h (p < 0.001), with Chromogranin A accumulating apically, toward the outer segment (Fig. 5j).

At the receptor level, murine OSM signals almost exclusively through the OSMR-gp130 heterodimer ^97^. Given the lack of *Osmr* expression in photoreceptors and the apparent delay in their response, the photoreceptor phenotype may arise via Müller glia-derived paracrine signals rather than direct OSM action. Receptor-ligand analysis shows that OSM-induced reactive glia upregulate a broad array of paracrine ligands (e.g., *Fgf2, Igf2*, and *Tnc*) targeting photoreceptor receptors (Extended Data Fig. 9d). Additionally, the downregulation of essential Müller glia genes (*Rlbp1, Glul*) (Fig. 5b) points to a disruption of metabolic and visual cycle support to the photoreceptors. However, since murine OSM might retain affinity for the LIFR-gp130 complex ^97,98^, and given that photoreceptors express *Lifr* (Extended Data Fig. 7a), OSM could also signal directly through LIFR-gp130. We exposed organoids to LIF, a LIFR-gp130-specific agonist, and found that LIF significantly induced both Müller glia reactivity and an increased proportion of ATF3^+^ photoreceptors (p < 0.01), confirming that LIFR-gp130 activation alone is sufficient to recapitulate the phenotype (Extended Data Fig. 9e).

Activation of the gp130 receptor complex, whether dimerized with OSMR or LIFR, recruits multiple downstream signaling cascades, including the JAK/STAT, NF-*κ*B, PI3K/Akt, and MAPK pathways ^94^. To isolate the specific cascades responsible for the OSM-induced phenotype, we co-treated organoids with OSM and inhibitors of these pathways. In particular, given the observed phosphorylation of STAT3 and the selective upregulation of its downstream targets (Fig. 5b,c,e,f, Extended Data Fig. 9a), we selected TPCA-1, a potent inhibitor of IKK-2 (NF-*κ*B) activity and STAT3 phosphorylation/transactivation ^99^, and Ruxolitinib, a selective JAK1/2 inhibitor ^100^. Co-treatment with either inhibitor completely prevented OSM-induced STAT3 phosphorylation and GFAP induction in Müller glia (non-significant vs control) (Fig. 5i). Both Ruxolitinib and TPCA-1 further prevented the OSM-induced apical CgA accumulation in photoreceptors (non-significant vs control) (Fig. 5j).

To determine whether these reactive macroglial programs are shared across the CNS, we perturbed astrocytes in mouse brain organoids using similar interventions. OSM induced astrocyte reactivity, characterized by a significantly increased proportion of phospho-STAT3^+^, Serpin A3N^+^ and HspB1^+^ astrocyte nuclei (p < 0.001) (Extended Data Fig. 10a). In contrast, extended, high-dose treatment with TNF*α*+HBEGF failed to induce astrocyte reactivity, highlighting important signaling distinctions between retinal and brain macroglia (Extended Data Fig. 10a). Activation of the LIFR-gp130 signaling complex with LIF also triggered astrocyte reactivity (p < 0.001) (Extended Data Fig. 10a), while STAT3/NF-*κ*B inhibition via TPCA-1 prevented OSM-induced astrocyte reactivity (p > 0.05 vs. untreated controls) (Extended Data Fig. 10b), confirming a shared gp130-STAT3-mediated mechanism across CNS macroglia.

Together, these results demonstrate that certain inflammatory cytokines, notably OSM and TNF*α*+HBEGF, drive Müller glia into distinct reactive states and photoreceptors into a degeneration-like trajectory culminating in apoptosis. Additionally, this data nominates the gp130-STAT3 signaling axis as a targetable driver of macroglial reactivity and photoreceptor stress.

## Discussion

Here, we built a single-cell atlas of mouse neural organoids encompassing multiple CNS regions and demonstrated the reliable emergence of matured neurons and macroglia capable of interaction within five weeks. Benchmarking against in vivo mouse atlases confirmed the presence of post-natal transcriptomic states that current human neural organoid protocols struggle to recapitulate ^24,25^. We further validated this rapid maturation across multiple modalities, notably reporting the presence of perineuronal nets after 60 days, and the emergence of synchronized electrophysiological networks within 4–5 weeks, in contrast to 4–6 months in human neural organoids. Together, this places mouse neural organoids as modular, rapidly developing systems to explore matured mammalian brain physiology.

Disease-associated macroglial transcriptomic states have been found in various human and murine CNS pathologies ^6,101,102^. Single-cell transcriptomic profiling of mouse organoids showed that macroglia act as response hubs to CNS stress pathway modulation and can adopt specific reactive states. Importantly, we demonstrated that these *in vitro* states recapitulate in vivo transcriptomic signatures of CNS injury and disease ^6,79,80^, which opens avenues for translating in vivo models to *in vitro* settings.

To explore the Müller glia phenotypic space, we generated a large single-cell transcriptomic perturbation atlas targeting 13 Müller glia-specific pathways. This atlas notably revealed that the Notch inhibitor DAPT combined with the histone deacetylase Trichostatin A shifted Müller glia into a reprogramming-like state, complementing previous genetic studies ^86,87^ and suggesting pharmacological approaches to Müller glia reprogramming. In parallel, Activin/TGF-*β* inhibition via SB431542 biased photoreceptor progenitor cells toward a cone rather than a rod trajectory. Although the direct mechanistic link between Activin/TGF-*β* signaling and photoreceptor fate decision remains unresolved, these results open exciting research avenues into rod-cone specification.

Müller glia reactivity is a hallmark of certain retinal pathologies, such as autoimmune ^101^ or diabetic retinopathies ^103^, yet the balance between its neuroprotective and detrimental effects remains poorly understood ^8,104^. While both Oncostatin M (OSM) and TNF*α*+HBEGF have been shown to induce the classical glial reactivity markers GFAP or phospho-STAT3 ^84,85,105,106^, our high-information, single-cell resolution profiling reveals that these cytokines induce distinct reactive Müller glia states. In particular, the OSM-induced state was uniquely characterized by upregulation of a chaperone-associated proteostatic program including *Hspb1, Cryab*, and *Clu*. Given the neuroprotective capacity of HspB1 and Clusterin ^107,108^, this suggests that Müller glia exist in different states with varying protective potential that could be harnessed therapeutically. Interestingly, OSM but not TNF*α*+HBEGF is sufficient to trigger reactivity in brain astrocytes, highlighting crucial differences in macroglial programs across organs. Further studies to resolve the molecular and regulatory mechanisms driving the distinct macroglial reactivity programs across organs and stimuli are warranted. In particular, as activated microglia are a primary physiological source of OSM ^109,110^, future iterations of mouse neural organoids incorporating microglia co-cultures will be essential to understand their role in retinal inflammation.

Photoreceptor stress and degeneration is multifactorial and often involves loss of proteostasis ^111^. OSM and inflammatory cytokines drove photoreceptors into distinct states, characterized by upregulation of gene programs including ER stress, and increased levels of ATF3 and Chromogranin A ^112^. Although the precise role of Chromogranin A in the retina remains largely unexplored, its upregulation in stressed photoreceptors and its implication in brain degeneration ^113,114^ suggests a previously undescribed role in photoreceptor stress response. When benchmarked against an in vivo degeneration trajectory from the rd10 mouse model ^93^, the ‘Stressed III’ state exhibited the strongest transcriptomic overlap, nominating it as a degeneration-like state driven by STAT3 signaling. Activation of gp130 signaling via OSM or LIF induced Müller glia reactivity and photoreceptor ER stress. Because photoreceptors express *Lifr* but not *Osmr* (Extended Data Fig. 7a), we propose that the OSM-induced photoreceptor phenotype is either mediated by macroglia, a direct result of LIFR-gp130 activation in photoreceptors, or a combination of both. Inhibiting the STAT3 axis with TPCA-1 or Ruxolitinib prevented the OSM-induced Müller glia and photoreceptor phenotypes, establishing the gp130-STAT3 axis as the driver of these perturbed states. Similar results in brain astrocytes further suggest this axis is shared and targetable across the CNS. Future studies utilizing conditional knock-outs of cytokine receptors and dense temporal profiling will be essential to disentangle the direct versus indirect influence of reactive Müller glia on photoreceptor health, and map the trajectory relationships between these stressed states and eventual photoreceptor death.

Altogether, this study establishes mouse neural organoids as a scalable, rapidly maturing platform to dissect neuro-macroglial biology, providing unprecedented access to macroglial phenotypic space and neuronal stress in the CNS.

## Limitations of the study

While human and mouse neural organoids present a valuable tool for neurobiology, they remain reductionist models with structural and physiological constraints. These include the absence of critical cell-cell interactions (e.g., with microglia, retinal pigment epithelium, and long-range innervating tissues), sensory inputs, extrinsic biomechanical forces, and vasculature, which altogether caps cellular maturation and causes the loss of specific cell types (e.g. retinal ganglion cells) over time. Our experimental design and profiling modalities also carry specific caveats. For instance, bulk metabolomics cannot distinguish intrinsic cellular metabolic shifts from within-organoid spatial gradients or changes in cell-type composition. Although macroglial EAATs mediate the majority of CNS glutamate clearance, L-trans-2,4-PDC also inhibits photoreceptor EAATs, which may contribute directly to the photoreceptor phenotype. Additionally, our perturbation screens utilized relatively short exposure timescales (12 hours to 4 days), and while this captures acute macroglial reactivity and the onset of cellular distress, it does not fully model prolonged progression of neurodegeneration. Furthermore, while we corroborated core mechanistic nodes at the protein level, much of our phenotypic analysis relies on transcriptomic inference, and cannot therefore account for complex behaviors like ECM remodeling and post-translational regulation. Finally, pharmacological interventions indiscriminately affect all cell types, preventing the definitive uncoupling of direct cytokine effects from indirect, macroglia-mediated effects. Looking ahead, mouse neural organoids could be a critical tool to resolve some of these limitations, given their potential for iterative bioengineering (e.g. incorporating missing cell types, vascularization, multi-region interactions, scaffolded cytoarchitecture, etc.) and benchmarking against in vivo counterparts.

## Acknowledgements

We are grateful to Barbara Treutlein, Anissa Kempf and Caren Norden for their supervision and valuable scientific insights, Sofia Tsepia for supporting pathology processes, and the Genomics 360 Lab of Roche Pharma Research and Early Development for their assistance with sequencing. We also thank Zhisong He and Jonas Simon Fleck for fruitful discussions. M.Si. thanks the University of Basel Biozentrum International PhD Program for training and scientific exchanges. J.-S.D. acknowledges support from the Doctoral Programme in Translational Engineering of Biological Systems established between ETH Zurich and F. Hoffmann-La Roche Ltd.

## Author Contributions

J.-S.D., M.Si., J.R., M.A.-B. and J.S. cultured, dissociated and processed mouse neural organoids for the pan-region scRNA-seq atlas. M.Si. and J.-S.D. cultured the organoids used in the rest of this study. M.Si. and J.-S.D. performed all perturbation experiments. J.-S.D. and M.Si. dissociated and processed organoids for scRNA-seq for the rest of this study. L.G.-T., M.A.-B., and M.Si. generated sequencing libraries. L.A. and J.-S.D. performed single-cell analysis quality control and phylomorphospace analysis. J.-S.D. and M.Si. performed clustering, annotations, and *in vivo* signature scoring. L.A. performed reference atlas integration, presence score estimation, receptor-ligand analysis, differential expression amplitude and correlation analyses, cell type proportion testing with inputs from J.-S.D. and M.Si.; J.-S.D. performed gene set enrichment analysis. M.Si. performed cell fate transition analysis. M.Si., J.-S.D. and I.C. performed immunofluorescence staining. M.Si. and J.-S.D. performed analysis of immunofluorescence data. J.-S.D., L.S., J.R., A.H., M.Sc. and J.G.C. designed the electrophysiological study. L.S., J.-S.D., T.P.B. and M.Sc. cultured organoids for electrophysiological measurements. L.S., T.P.B. and M.Sc. performed measurements and analysis of electrophysiological data. J.-S.D., T.F., M.Z. and J.G.C. designed the metabolomics study. T.F. performed mass spectrometry and data pre-processing. J.-S.D. analyzed the metabolomics data. L.A., J.R., M.L., and J.G.C. provided supervision and support. J.-S.D., M.Si. and J.G.C. designed the study and wrote the manuscript. All authors read and approved the manuscript.

## Competing Financial Interests

All authors associated with the Roche Institute of Human Biology are employees of Hoffmann-La Roche AG. The company provided support in the form of salaries for these authors but did not have any additional role in the study design, data collection and analysis, decision to publish or preparation of the manuscript. The other authors declare no conflict of interest.

## Data Availability

All sequencing, image analysis, metabolomic, electrophysiological data and code used in this study will be made publicly available via GitHub, Zenodo and the Gene Expression Omnibus upon publication. Please contact the corresponding authors for inquiries.

## Declaration of AI-Assisted Technologies in the Writing Process

During the preparation of this work the authors used large-language models (Google Gemini v3.0 Pro, Claude Sonnet 4.5) for language editing and stylistic refinement. The authors reviewed, edited, and approved all AI-generated suggestions and take full responsibility for the content of the published article.

## Methods

### Mouse embryonic stem cell culture

The murine ESC line SBr: CGR8(Sox1eGFP;BramCherry) ^115^ has been generously provided to the Roche Institute for Human Biology by David Suter. The mESC were cultured in mESC medium (DMEM high glucose + GlutaMax (Gibco #61965026) supplemented with 10% v/v ES-qualified FBS (Thermo Fisher #16141079), 1X MEM-NEAA (Gibco #11140035), 1X sodium pyruvate (Gibco #11360039), 0.1 mM *β*-mercaptoethanol (Gibco #31350010), 1X Penicillin-streptomycin (Gibco #15140122)) supplemented with 2i (3 *µ*M CHIR99021 (StemCell technologies #72054) + 1 *µ*M PD0325901 (Selleck Chemicals #S1036)) and 100 *µ*g/ml LIF (ESGRO Mouse LIF, Sigma #ESG1106) in a humidified incubator (5% CO_2_, 37°C). The cells were split every other day, by washing with PBS (Gibco #10010056) and dissociating with Accutase (StemCell #07920) for 3 min at room temperature. The cell suspension was collected in a tube and centrifuged at 300g for 3 min. After centrifugation, the cell pellet was resuspended, counted and seeded with a density of 50000 cells per well of a six-well plate (CytoOne #CC7682-7506) in 2 ml of mESC medium.

### Mouse neural organoid culture

#### Mouse minimally guided brain organoids

Mouse minimally guided brain organoids (referred to as brain organoids) were derived following an adapted version of a human minimally guided protocol ^20,116^ using the murine ESC line SBR. Briefly, mouse ES cells (passage <20) were dissociated with Accutase (StemCell #07920) and 3000 cells were plated in each well of an ultra-low-attachment 96-well plate (Sbio #MS9096UZ or Corning #4520) in modified hES Medium (4:1 DMEM-F12 (Gibco #11330032) and KnockOut Serum Replacement (Gibco #10828028) supplemented with 3% v/v ESC-qualified FBS (Gibco #16141079), 1X GlutaMAX (Gibco #35050061), 1X MEM-NEAA (Gibco #11140035), 1X Penicillin-streptomycin (Gibco #15140122), 0.1 mM 2-mercaptoethanol (Gibco #31350010) and 4 ng/mL of murine FGF2 (Peprotech #450-33)). On day 2 and 4, media was exchanged for Neural Induction Medium (NIM: DMEM-F12 (Gibco #11330032) supplemented with 1X N2 supplement (Gibco #17502048), 1X GlutaMAX (Gibco #35050061), 1X MEM-NEAA (Gibco #11140035), 1X Penicillin-streptomycin (Gibco #15140122) and 1 *µ*g/mL of Heparin (Sigma #H3149)). On day 5, media was replaced by Differentiation Media minus Vitamin A (IDM-A: 1:1 DMEM-F12 (Gibco #11330032) and Neurobasal (Gibco #21103049) supplemented with 0.5X N2 supplement (Gibco #17502048), 1X B27 supplement without Vitamin A (Gibco #12587010), 1X GlutaMAX (Gibco #35050061), 0.5X MEM-NEAA (Gibco #11140035), 0.1 mM 2-mercaptoethanol (Gibco #31350010), 1X Penicillin-streptomycin (Gibco #15140122) and 5 *µ*g/mL human insulin (Sigma #I9278)) supplemented with 2.5% v/v Matrigel (Corning #354234). On day 7, organoids were washed three times in cold Differentiation Medium minus Vitamin A to remove Matrigel and transferred (32 organoids per dish) to 90 mm ultra-low-attachment cell culture dishes (Sbio #MS-90900) on an orbital shaker (set to 63 rpm). From day 8 onwards, organoids were maintained in Differentiation Medium with Vitamin A (IDM+A: 1:1 DMEM-F12 (Gibco #11330032) and Neurobasal (Gibco #21103049) supplemented with 0.5X N2 supplement (Gibco #17502048), 1X B27 supplement with Vitamin A (Gibco #17504044), 1X GlutaMAX (Gibco #35050061), 0.5X MEM-NEAA (Gibco #11140035), 0.1 mM 2-mercaptoethanol (Gibco #31350010), 1X Penicillin-streptomycin (Gibco #15140122), 5 *µ*g/mL human insulin (Sigma #I9278), 0.4 mM Ascorbic acid (Sigma #A4544) and 1 mg/mL Sodium bicarbonate (Sigma #S5761)), with media changes every other day.

For the automated version of this protocol, organoids were started from 500 cells, an ideal balance between minimizing apoptosis at the organoid core and maximizing reproducibility in organoid morphology (Extended Data Fig. 1b), and maintained in ultra-low-attachment 96-well plates (Sbio #MS-9096UZ or Corning #4520) until the experimental end point. Organoids were exposed to Matrigel as 1–3.5% v/v Matrigel in solution and washed using *≥*3 consecutive half media change with cold medium on day 7. After day 8, medium was exchanged more frequently (every 24–48h) to account for the lack of shaking.

#### Mouse cortical organoids

Mouse cortical organoids were derived following an adapted version of Velasco et al. ^39^ using the murine ESC line SBR. Briefly, mouse ES cells (passage 20) were dissociated by Accutase treatment (StemCell #07920) and 9000 cells were plated in each well of an ultra-low-attachment 96-well plate (Sbio #MS-9096UZ) in CDM1 medium (GMEM (Gibco #11710035) supplemented with 20% v/v KnockOut Serum Replacement (Gibco #10828028), 1 mM Sodium Pyruvate (Gibco #11360039), 1X MEM-NEAA (Gibco #11140035), 1X Penicillin-streptomycin (Gibco #15140122), 0.1 mM *β*-mercaptoethanol (Gibco #31350010), 3 *µ*M IWR1 (Selleckchem #7086) and 5 *µ*M SB431542 (Selleckchem #S1067)). On day 3 and 5, medium was exchanged for fresh CDM1 medium. On day 7, organoids were transferred (32 organoids per dish) to 90 mm ultra-low-attachment cell culture dishes (Sbio #MS-90900) on an orbital shaker (set to 63 rpm) and medium was replaced with CDM2 medium (DMEM-F12 (Gibco #11330057) supplemented with 1X N2 supplement (Gibco #17502048), 1X Chemically Defined Lipid Concentrate (Gibco #11905031) and 1X Penicillin-streptomycin (Gibco #15140122)). On day 10 and 13, medium was exchanged for CDM3 medium (DMEM-F12 (Gibco #11330057) supplemented with 1X N2 supplement (Gibco #17502048), 1X Chemically Defined Lipid Concentrate (Gibco #11905031), 1X Penicillin-streptomycin (Gibco #15140122), 10% v/v ESC-qualified FBS (Gibco #16141079), 5 *µ*g/mL Heparin (Sigma #H3149-25KU) and 1% v/v Matrigel (Corning #354234)). Finally, from day 16 onwards, medium was exchanged every other day with CDM4 medium (DMEM-F12 (Gibco #11330057) supplemented with 1X N2 supplement (Gibco #17502048), 1X B27 supplement (Gibco #17504044), 1X Chemically Defined Lipid Concentrate (Gibco #11905031), 1X Penicillin-streptomycin (Gibco #15140122), 10% v/v ESC-qualified FBS (Gibco #16141079), 5 *µ*g/mL Heparin (Sigma #H3149-25KU) and 2% v/v Matrigel (Corning #354234).

#### Mouse thalamic organoids

Mouse thalamic organoids were derived following an adapted version of Shiraishi et al. ^40^ using the murine ESC line SBR. Briefly, mouse ES cells (passage20) were dissociated by Accutase treatment (StemCell #07920) and 500 cells were plated in each well of an ultra-low-attachment 96-well plate (Corning #4520) in gfCDM medium (1:1 IMDM (Gibco #21980032) and F-12 (Gibco #21765029) supplemented with 1% v/v Chemically Defined Lipid Concentrate (Gibco #11905031), 1X Penicillin-streptomycin (Gibco #15140122), 450 *µ*M Monothioglycerol (Sigma #M1753), 15 *µ*g/mL murine apo-transferrin (Sigma #T0523), 5 mg/mL BSA (Sigma #A4919)) supplemented with 10 *µ*M SB431542 (Selleckchem #S1067) and 1 *µ*g/mL human insulin (Sigma #I9278). On day 2 the medium was exchanged for fresh gfCDM medium supplemented with 10 *µ*M SB431542 and 1 *µ*g/mL human insulin. On day 4 the medium was exchanged for gfCDM medium supplemented with 10 *µ*M SB431542, 1 *µ*g/mL human insulin, 1 *µ*M PD0325901 (Selleckchem #S1036) and 30 ng/mL human BMP7 (Peprotech #120-03P). On day 7 and 9 the medium was exchanged for Neural Induction Medium (NIM, see Mouse minimally guided brain organoids section). From day 12 onwards, medium was exchanged 1–2 days with fresh Differentiation Media with Vitamin A (IDM+A, see Mouse minimally guided brain organoids section).

#### Mouse midbrain organoids

Mouse midbrain organoids were derived following a custom protocol inspired by Qian et al. ^41^ and Reumann et al. ^117^ using the murine ESC line SBR. Briefly, mouse ES cells (passage <25) were dissociated by Accutase treatment (StemCell #07920) and 500 cells were plated in each well of an ultra-low-attachment 96-well plate (Corning #4520) in modified hES Medium (see Mouse minimally guided brain organoid section). On day 1, medium was replaced with modified hES Medium supplemented with 1.5 *µ*M CHIR99021 (StemCell Technologies #72054). On day 2 and 4, medium was exchanged for Neural Induction Medium (NIM, see Mouse minimally guided brain organoid section) supplemented with 1.5 *µ*M CHIR99021 (StemCell Technologies #72054), 300 nM SAG (StemCell Technologies #73412) and 100 ng/mL murine FGF8b (Peprotech #100-25). On day 5, medium was replaced with Differentiation Medium minus Vitamin A (IDM-A, see Mouse minimally guided brain organoids section) supplemented with 1.5 *µ*M CHIR99021 (StemCell Technologies #72054), 300 nM SAG (StemCell Technologies #73412) and 100 ng/mL murine FGF8b (Peprotech #100-25). On day 6, medium was replaced with non-supplemented Differentiation Medium minus Vitamin A. From day 8 onwards, medium was exchanged every 1–2 days with fresh Differentiation Media with Vitamin A (IDM+A, see Mouse minimally guided brain organoids section).

#### Mouse cerebellum organoids

Mouse cerebellum organoids were derived following an adapted version of Muguruma et al. ^42^ using the murine ESC line SBR. Briefly, mouse ES cells (passage < 20) were dissociated by Accutase treatment (StemCell #07920) and 500 cells were plated in each well of an ultra-low-attachment 96-well plate (Corning #4520) in gfCDM medium (see Mouse thalamic organoid section) supplemented with 7 *µ*g/mL human insulin (Sigma #I9278). On day 1, 3 and 5 medium was exchanged with gfCDM medium supplemented with 7 *µ*g/mL human insulin (Sigma #I9278) and 20 ng/mL murine FGF2 (Peprotech #450-33). On day 7 and 9, medium was replaced with gfCDM medium supplemented with 7 *µ*g/mL human insulin (Sigma #I9278), 20 ng/mL murine FGF2 (Peprotech #450-33), 1 nM human BMP4 (Peprotech #120-05ET), 10 *µ*M Cyclopamine (StemCell Technologies #72074) and 30 nM murine SHH (Peprotech #315-22). On day 11 and 13, medium was replaced with Neural Induction Medium (NIM, see Mouse minimally guided brain organoids section). From day 15 onwards, medium was exchanged every 1–2 days with fresh Differentiation Media with Vitamin A (IDM+A, see Mouse minimally guided brain organoids section).

#### Mouse retinal organoid

Mouse retinal organoids were generated from mouse ESC based on a protocol from Decembrini et al. ^43^ with some modifications inspired by the trisection approach from Völkner et al. ^44^. Briefly, mouse ES cells (passage <20) were dissociated by Accutase treatment (StemCell #07920) and 3000 cells were plated in each well of an ultra-low-attachment 96-well plate (S-bio #MS-9096UZ) in 120 *µ*L of medium. The media for the first 24h is 50% mESC medium without 2iLIF and 50% Retinal Differentiation Medium (RDM: GMEM (Gibco #11710035), 1.5x KnockOut™ Serum Replacement (Gibco #10828028), 1X MEM-NEAA (Gibco #11140035), 1x sodium pyruvate (Gibco #11360039), 0.1 mM *β*-mercaptoethanol (Gibco #31350010), 1X Penicillin-streptomycin (Gibco #15140122), 1X N2 (Gibco #17502-048), 1X B27 w/o VitA (Gibco #12587-010)). On day 1, day 3 and day 5, a half media change with matrigel (Corning® Matrigel® Growth Factor Reduced #356231) was performed to reach a final concentration of 2% of matrigel in the media. On day 7 the organoids were collected from the 96WP using a wide-bore tip (ThermoFisher #2079GPK) and 48 organoids were collected in one falcon tube. To remove the matrigel, the organoids were washed three times with ice cold media (DMEM/F-12 +GlutaMAX (Gibco #31331-028). Next, 48 organoids were transferred into a 9cm dish (MS-9090XZ PrimeSurface) with 10ml of Retinal Maturation Medium (RMM: DMEM/F-12 +GlutaMAX (Gibco #31331-028), 1X N2 (Gibco #17502-048), 1X Penicillin-streptomycin (Gibco #15140122)) and placed in an incubator on an orbital shaker (set to 63 rpm). On day 9 a full media change (12ml) was performed with the RMM supplemented with 1x B27 with VitA (Gibco #12587001). On day 12 a media change was performed with 15ml of the RMM supplemented with 0.5 *µ*M 9-cis Retinoic Acid (Sigma #R5754) and 1mM Taurine (Sigma #T8691) (RMM+RA+Tau). On day 14 or 15, 24 organoids were transferred to a new 9cm dish with 2ml of media, the dish was tilted and with a P1000 pipette the organoids were dissociated to small epithelial fragments. Then the medium was changed with 15ml RMM +RA +Tau. The pieces recover and form closed organoid structures after 24h. This step will lead to an increased yield of epithelium and to smaller organoids with less of a necrotic core. From day 16 onwards the media was changed three times a week with 20ml of RMM +RA +Tau per dish and kept on an orbital shaker. For treatments, 10–15 organoids were transferred into one well of a 6 well plate (Corning #3471) in 3ml of media. The organoids were manually picked by visual assessment using a brightfield microscope (Invitrogen™ EVOS™ XL Core Imaging System). Organoids that did not display a shiny epithelium were not included into further experiments.

### Protocol automation, bright-field imaging and morphological analysis

The liquid handling and imaging for certain neural organoid protocols was performed on automated and semi-automated setups. Media changes during the differentiation phase of the protocols were performed on either a Washer Dispenser (Agilent BioTek EL406) or a programmable 96-manifold pipetting platform (Integra VIAFLO 96) under sterile conditions. Long-term maintenance was conducted using a custom-integrated liquid handling system comprising a programmable incubator, a refrigerated media storage unit connected via a programmable peristaltic pump, and a liquid handling platform (Tecan Freedom EVO 200). The workflow consisted in transferring media from cold storage to a sterile reservoir for temperature equilibration prior to performing automated media changes on plates retrieved from the programmable incubator. Automated bright-field imaging was performed on a custom-integrated plate handling system comprising a programmable incubator connected to a plate scanner (Molecular Devices ImageXpress Confocal HT.ai). For the optimization of seeding density (Extended Data Fig. 1c), mouse minimally guided brain organoids were started from 500 and 100 stem cells, fixed on day 20 (4% PFA for 24h at 4°C), imaged under bright-field on a spinning-disk confocal microscope (Nikon Ti2) at 10x resolution, and image tiles were stitched together using the Nikon NIS-Elements software. For morphological tracking along organoid development (Extended Data Fig. 1d), bright-field images were automatically taken every 24h on the platform described above at 10x magnification. Generally, tiled images were masked, and morphological metrics extracted using python’s skimage and scipy packages. All other bright-field images in this study were acquired on a benchtop bright-field microscope (Olympus CKX53) at 4x resolution unless otherwise mentioned.

### Single-cell dissociation and RNA sequencing library preparation

Organoid dissociation for scRNA-seq was performed using the Neural Tissue Dissociation Kit (P) (Miltenyi Biotec #130-092-628). All tubes and pipette tips used during the procedure were either of commercially available low-binding types (e.g., Eppendorf LoBind© range) or pre-coated with DPBS −Ca^2+^/−Mg^2+^ (Gibco #14190250) supplemented with 1% v/v MACS BSA (#130-091-376) to prevent cellular adhesion on the plastic. Large organoids (*>~* 2mm in diameter) were first cut into small fragments using a scalpel in a Petri dish filled with DPBS −Ca^2+^/−Mg^2+^ (Gibco #14190250). 1–8 organoids replicate from a single condition were then transferred into a 5 mL conical tube using a wide-bore pipette tip (ThermoFisher #2079GPK), allowed to settle in DPBS −Ca^2+^/−Mg^2+^ (Gibco #14190250), and the supernatant was removed. For each dissociation reaction, 1950 *µ*L of pre-warmed Papain mix (50 *µ*L of Enzyme P in 1900 *µ*L of Buffer X) was added and the tube was incubated for 5–10 minutes at 37°C. For more mature or complex samples (e.g. brain organoids older than 20 days), all incubation steps were performed on an orbital shaker set to 68 rpm. After this first incubation, 90 *µ*L of DNAse mix (30 *µ*L of Enzyme A in 60 *µ*L of Buffer Y) was added and mixed in by gently inverting the tube. The sample then underwent a series of gentle mechanical dissociations using a P1000 pipette, initially with a wide-bore tip, followed by a standard tip, interspersed with 5- to 10-minute shaking incubations at 37°C. This was repeated for a total of 15–50 minutes until a single-cell suspension was visually confirmed. The single-cell solution was then filtered 1–2x through a 30 or 70 *µ*m strainer (Miltenyi Biotec #130-095-823 #130-095-823) pre-wetted with ice-cold DPBS −Ca^2+^/−Mg^2+^ (Gibco #14190250) containing 0.5% BSA, followed by a 2 mL wash with the same buffer. Cells were pelleted by centrifugation at 300 x g for 3 minutes at 4°C and gently resuspended in 100–500 *µ*L of cold DPBS with 0.5% BSA. Cell concentration and viability were assessed using Trypan Blue (Invitrogen #T10282) using an automated cytometer (Thermo Fisher Scientific, Countess II or Countess 3 FL). The single-cell suspension was subsequently diluted to the optimal concentration, loaded onto the 10x Genomics microfluidic chip with a target recovery of 10,000 cells, and single-cell libraries were prepared using the Next GEM Single Cell 3^*′*^ v3.1 kit according to manufacturer instructions.

For the multiplexed single-cell RNA-seq screen using the Parse Biosciences pipeline, dissociation was adapted for a 96-well format. 8 retinal organoids per independent replicate were transferred into a 96-well 1 mL U-bottom deep-well plate (Eppendorf #0030501241) containing DPBS −/−. Organoids were enzymatically digested by adding 500 *µ*L of a 2X Papain mix (25 *µ*L Papain in 500 *µ*L buffer X) per well, followed 15 minutes later by 15 *µ*L of the DNAse mix (10*µ*L buffer Y + 5*µ*L DNAse). The tissue was incubated at 37°C for a total of 60 minutes. Mechanical dissociation was performed at 10- to 15-minute intervals using harsh multichannel pipetting. Immediately prior to filtration, samples were vigorously triturated 12 times with a P200 pipette followed by 8 times with a P1250 pipette. The resulting cell suspensions were passed through a 30 *µ*m 96-well plate strainer (CS CRIE #R-CSC001-01A) into a 96-well 0.8 mL V-bottom deep-well plate (Abgene #AB0765). Cells were pelleted at 300 x g for 3 minutes at 4°C, washed in 650 *µ*L of cold DPBS −Ca^2+^/−Mg^2+^ containing 0.5% BSA, and spun again at 200 x g for 2 minutes to remove small debris. Finally, cells were resuspended in 600 *µ*L of cold DPBS −Ca^2+^/−Mg^2+^ with 0.5% BSA. Following cell counting, volumes were adjusted to normalize the sample to approximately 750,000 cells per well before proceeding to cell fixation and permeabilization using the Evercode Cell Fixation v3 High-Throughput Plate Workflow kit (Parse Biosciences, User Manual UM0029 v1.1, #ECFC3501), with several modifications. The initial single-cell centrifugation was modified to 3 minutes at 200g, and the centrifugation following the addition of the Fix and Perm Stop Buffer was adjusted to 5 minutes at 300g. To prevent sample loss, all plate strainer steps standard to the protocol were omitted. The volume of Cell Storage Master Mix added during the final resuspension step was adjusted based on the initial cell input and residual volume to achieve a target minimum final concentration of 3000 cells/*µ*L, adding a maximum of 55 *µ*L of Storage Master Mix to the residual volume in each well. Downstream split-pool combinatorial barcoding and library preparation were subsequently performed with Evercode Mega v3 reagents (Parse Biosciences, User Manual UMWT3500, #ECWT3501) according to the manufacturer’s instructions.

### Single-cell RNA sequencing analysis

#### Raw data processing and quality control

Raw count matrices were created from sequencing results using CellRanger v7.1.0 with the option to include intronic reads and imported in R using the Seurat package. Initial quality control (QC) removed low-quality cells, empty droplets, and dying cells by applying sample-specific filtering thresholds for the minimum and maximum number of detected genes, total transcript counts, and maximum allowable percentage of mitochondrial transcripts. The DoubletFinder algorithm was utilized to estimate and filter out heterotypic multiplets. For the multiplexed perturbation dataset, a multi-modal consensus approach was developed to flag the large cellular aggregates created during the fixation step of the library preparation by intersecting DoubletFinder doublet classifications from scrublet with orthogonal detection methods which included transcript count bimodal fitting and mutually exclusive cell-type-marker entropy scoring. After data pre-processing and clustering, seldom clusters of doublet cells were identified in certain datasets based on their mixed expression profiles and filtered out. In the interferon perturbation dataset, two clusters of hindbrain neurons and oligodendrocytes only found in the Interferon treatment were deemed to reflect variability in brain region identity in the organoids pooled for sequencing rather than treatment-induced effects, and therefore excluded from the dataset before Differential Expression Analysis to prevent confounding the results.

#### Pre-processing, tissue atlas integration, and dimensionality reduction

Post-QC count matrices were log-normalized, and the most variable features (typically 2,000 to 3,000 genes) were selected for downstream modeling. The data was subsequently scaled, and Principal Component Analysis (PCA) was performed to reduce dataset dimensionality. To correct for batch effects, datasets were integrated using the Harmony algorithm applied over the principal components.

For the integrated retina atlas, the organoid dataset and the reference tissue were integrated individually with scVI ^118^, each trained with 2 hidden layers, a latent dimension of 30, negative binomial gene likelihood, and a random seed 0. On the resulting scVI latent representations, k-nearest neighbor graphs (*k* = 15) and UMAP embeddings were computed. The organoid dataset was clustered using the Leiden algorithm (resolution = 2.0) implemented in Scanpy ^119^. Both organoid and reference tissue clusters were annotated using Snapseed ^25^, leveraging a hierarchically organized set of canonical retinal cell type markers (2– 4 marker genes per cell type). To integrate the reference and organoid datasets, datasets were merged, and the top 2000 common highly variable genes were selected. Integration was performed using scANVI ^120^ by initializing the pretrained scVI model with a maximum of 20 epochs and 100 samples per label, using the cell type annotations from the previous step. Following scANVI integration, k-nearest neighbor graphs (*k* = 15) and UMAP embeddings were computed on the resulting scANVI latent representation. The cell type annotation was manually refined for the cell types uniquely found in the reference or the organoid dataset (fibroblasts, endothelial cells) and 1043 cells were identified as doublets based on the expression of shared marker genes of multiple cell types and removed before further analysis.

For the integrated brain atlas, both the reference tissue and the organoid datasets were integrated using scVI, utilizing 2 hidden layers, a latent dimension of 30, and a negative binomial gene likelihood. Next the organoid dataset was loaded into the pretrained scVI reference model. The query model was fine-tuned to adapt to the organoid-specific expression patterns, with a maximum of 200 epochs with weight decay set to 0.

#### Clustering and cell type annotation

Unsupervised clustering was carried out using Seurat’s graph-based clustering functions (FindNeighbors and FindClusters), tested across a range of resolutions to identify biologically coherent groupings. Clusters were annotated based on the expression of canonical marker genes and the genes uniquely expressed by each cluster (as found by FindAllMarkers). For higher-resolution annotation, key cell types (astrocytes, Müller glia, and photoreceptors) were computationally subsetted, independently scaled, iteratively re-clustered and annotated based on the expression of canonical marker genes and the genes uniquely expressed by each cluster (as found by FindAllMarkers). All other details can be found on the corresponding R, Markdown and Python scripts.

#### Presence score estimation

We estimated presence scores using an approach adapted from He et al. ^25^ via CellMapper v0.2.5 (https://zenodo.org/records/16606250). In brief, we constructed a weighted k-nearest neighbors (wKNN) graph based on the scVI latent embedding, utilizing the RAPIDS implementation (`method=‘rapids’`) with a neighborhood size of 500 (`neighbors=500`). Presence scores were then calculated using the `compute_presence_score` function (parameters: `groupby=‘batch’`, `log=True`, and `minmax=True`). Conceptually, for each reference cell subtype and organoid dataset, this subsets the wKNN graph to contain only the specific reference subtype alongside all cells from the organoid dataset, followed by aggregating the edges within this local neighborhood, effectively proxying the *in vitro* presence of this reference subtype. The resulting metrics were smoothed across the reference graph using a random-walk with restart algorithm to reduce noise with a hyperparameter *t* = 1000. Finally, we assigned each reference cell subtype a final presence score corresponding to the maximum smoothed score it achieved across all individual organoid datasets. A high presence score indicates that cells with similar molecular profiles are frequently and reliably observed in the organoid protocols.

#### Receptor-ligand analysis

For the *in vitro* versus *in vivo* comparison of receptor-ligand (R-L) pair expression (Extended Data Fig. 3a), interactions were modeled using the liana Python package. Prior to analysis, genes detected in fewer than 10 cells were excluded. R-L interactions were then inferred utilizing the `rank_aggregate` function against the mouseconsensus resource, with the minimum expression proportion (`expr_prop`) set to 0.1. The analysis was explicitly restricted to interactions between core cell types in the reference tissues (astrocytes, neurons, and oligodendrocytes for the brain; Müller glia, photoreceptors, and amacrine cells for the retina). Interacting pairs were strictly filtered for high specificity (specificity rank <0.1), and the top 50 pairs exhibiting the highest interaction strength (magnitude rank) were extracted for each distinct source-target cell type combination, resulting in approximately 100 genes per cell-type pair. Finally, to quantify cross-system fidelity, the average expression of these filtered genes was computed, and cosine similarity was calculated between corresponding cell types across the *in vivo* reference and *in vitro* organoid datasets. For the analysis of candidate Müller glia-photoreceptor ligand-receptor pairs in OSM-treated organoids (Extended Data Fig. 9d), reference LR pairs were retrieved from the liana R package’s MouseConsensus resource database. This list of LR pairs was filtered to include only pairs with ligands significantly upregulated in Müller glia in OSM-treated versus control organoids and receptors expressed in a minimum of 10% of rods or 10% of cones (and expressed in >20 cells) before being ranked by ligand log_2_ fold-change relative to control.

#### Compositional and transcriptomic similarity to reference tissues

To compare the cell type compositional profiles between the organoid dataset, the *in vivo* atlas, and the human organoid atlas, the cell type annotations were harmonized across the datasets of the same organs. To focus on major developmental lineages, a higher-level annotation was chosen, by pooling individual cell types into broader categories. For the reference datasets multiple timepoints were pooled to average variability in the data and to generate representative compositional profiles. Cell type proportions were calculated for each dataset and developmental stage. Cosine similarity was then calculated between the organoid and reference compositions for each developmental stage.

To enable label transfer of categorical annotations (e.g., developmental age) from reference tissues to organoid cells and facilitate comparison of developmental maturity, we employed CellMapper v0.2.5. CellMapper uses k-nearest neighbor matching within embedding spaces to transfer labels from the most transcriptomically similar *in vivo* tissue counterparts to query organoid cells. We applied scANVI embeddings for retinal organoids and scVI embeddings for brain organoids.

#### Differential expression analysis

For differential expression analysis, cell populations of interest were selected prior to pseudobulk aggregation. In the retina dataset, astrocytes and starburst amacrine cells were excluded due to their low representation across samples. In the brain dataset, analyses were restricted to neurons, astrocytes, oligodendrocyte precursor cells (OPCs), oligodendrocytes, intermediate progenitor cells (IPCs), and neural progenitor cells (NPCs). Lowly expressed genes were removed using a quantile-based filtering approach, retaining genes above the 20th percentile expression threshold in the retina dataset and above the 50th percentile threshold in the brain dataset. For the brain dataset, mitochondrial and ribosomal genes were additionally excluded. The remaining single-cell RNA-seq profiles were aggregated into pseudobulk samples by summing counts across cells belonging to the same sample and cell type. Library size normalization was performed by estimating sample-specific size factors using the ratio method. Differential expression was then assessed separately for each cell type using the delnx framework (v.0.2.2) (https://github.com/mlbelab/delnx). Gene-wise dispersions were estimated using a negative binomial model while accounting for both batch and condition effects. Differential expression testing was subsequently performed using a negative binomial generalized linear model, comparing each experimental condition against the control group within each cell type. Batch was included as a covariate during dispersion estimation to account for technical variation. Results from all cell-type-specific analyses were combined for downstream analyses and interpretation.

For the comparison between reactive Müller glia clusters we used the DESeq2 package for pseudobulking and differential testing, combined with adaptive shrinkage using the ashr package (https://github.com/stephens999/ashr) to estimate conservative log-fold changes.

#### Transcriptomic response amplitude and correlation analysis

To quantify perturbation responses across cell types, single-cell expression profiles were aggregated at the sample level within each cell type and perturbation condition. Cell type-perturbation groups represented by fewer than 10 cells were excluded prior to aggregation. Perturbation effect magnitude was defined as the average transcriptional distance between perturbed and control samples, measured using Euclidean distance in gene expression space. For each cell type, the mean expression profile of control samples was used as a reference.

To assess the extent to which perturbation responses were shared across cell types, perturbation-induced expression changes were computed as the difference between the mean perturbed and mean control expression profiles for each cell type. Pairwise Pearson correlations of these response vectors were then calculated across cell types for each perturbation. Perturbations represented in fewer than two cell types were excluded from this analysis. The average correlation of a given cell type with all other cell types was used as a measure of perturbation effect sharedness, with higher values indicating more broadly conserved transcriptional responses and lower values indicating greater cell-type specificity.

#### Gene set enrichment analysis

Gene Set Enrichment Analysis (GSEA) was performed on a per-cell-type basis by filtering differential expression results for each respective cell type. Genes were subsequently ranked by their log-fold change (or delnx coefficient). Enrichment was computed using the clusterProfiler package (v.4.0 or similar) against a combined biological database comprising the KEGG, WikiPathways, Reactome ontological gene sets, or a combination of them retrieved via the msigdbr R package (*Mus musculus*, Category C2), and configured with a minimum gene set size of 10 to 15 genes, a maximum size of 500 genes, and significance adjusted via Benjamini-Hochberg FDR correction. To eliminate redundant pathways and improve interpretability, a custom semantic filtering pipeline was implemented. This involved initially collapsing exact gene set duplicates, followed by the application of a Jaccard similarity filter (threshold = 0.80 to 0.95 depending on the dataset) to iteratively remove less significant terms that exhibited extensive gene set overlap with more significant ontology terms. For the multiplexed perturbation dataset, we used a custom filtering to extract “strongly enriched and treatment-specific” pathways: we scored candidate pathways by integrating their normalized enrichment score (NES) and statistical significance, while further enforcing a minimum Gini inequality threshold (e.g., *>* 0.2 to 0.45) across treatment conditions.

#### In vivo signature scoring

Signature gene sets were derived as described in Supplementary Table 5. Scoring was executed using either the AddModuleScore function from the Seurat package or the AddModuleScore_UCell from the UCell package. Effect size (Cohen’s d) was computed using the effsize package.

#### Cell type proportion change analysis

Changes in cell-type composition were assessed using the scanpro framework ^121^ (v0.4.1). For each experimental condition, the dataset was subsetted to include only cells from the condition of interest and the corresponding control samples. Cell types represented by fewer than 50 cells across the subsetted dataset were excluded from the analysis. Differential abundance testing was then performed using scanpro with a logit transformation of cell-type proportions (`transform = “logit”`), while all other parameters were kept at their default settings.

#### Phylomorphospace representation

The phylomorphospace representation for each primary cell type (Müller glia, Rods, Cones) was computed based on the Euclidean distance between the response expression vectors of each treatment. Multidimensional scaling (MDS) dimensionality reduction was performed via the base R cmdscale function. Simultaneously, a neighbor-joining (NJ) tree was constructed from the same Euclidean distance matrix utilizing the ape package. To integrate these metrics, the NJ tree was projected onto the 2D MDS embedding using the phylomorphospace function from the phytools package. Finally, continuous variables such as specific target gene expression, pathway signature scores, and the fraction of transcriptionally responding cells were overlaid onto the nodes of the phylomorphospace network. In this phylomorphospace representation, distances between dots reflect divergence in the transcriptomic response between the corresponding perturbations, while the NJ topology (i.e. how many branches away) reflects the hierarchical similarity of responses. Perturbations yielding responses similar in amplitude and direction correspondingly cluster together in the MDS plot and are connected by adjacent branches.

#### Cell fate transition probability analysis

To evaluate changes in cell fate transition probability upon SB431542 perturbation, the dataset was subsetted to rods, cones and photoreceptor progenitors from the control and the SB431542-treated condition. After preprocessing the resulting dataset as described above, the two conditions were integrated using Harmony (harmonypy, v2.0.0), and a neighbourhood graph was constructed (*k* = 50). CellRank2^122^ was used to calculate the fate probabilities. First, a transition matrix was computed using a combination of the PseudotimeKernel and the Connectivitykernel (0.8 *\* K*_pseudotime_ + 0.2 *\* K*_connectivity_). Next we used the GPCCA estimator to compute three macrostates (which corresponded to rods, cones and photoreceptor progenitors), set the ‘rods’ and ‘cones’ states as terminal states, and computed fate transition probabilities to these states for both control and SB431542 photoreceptor progenitors.

### List of antibodies used in this study

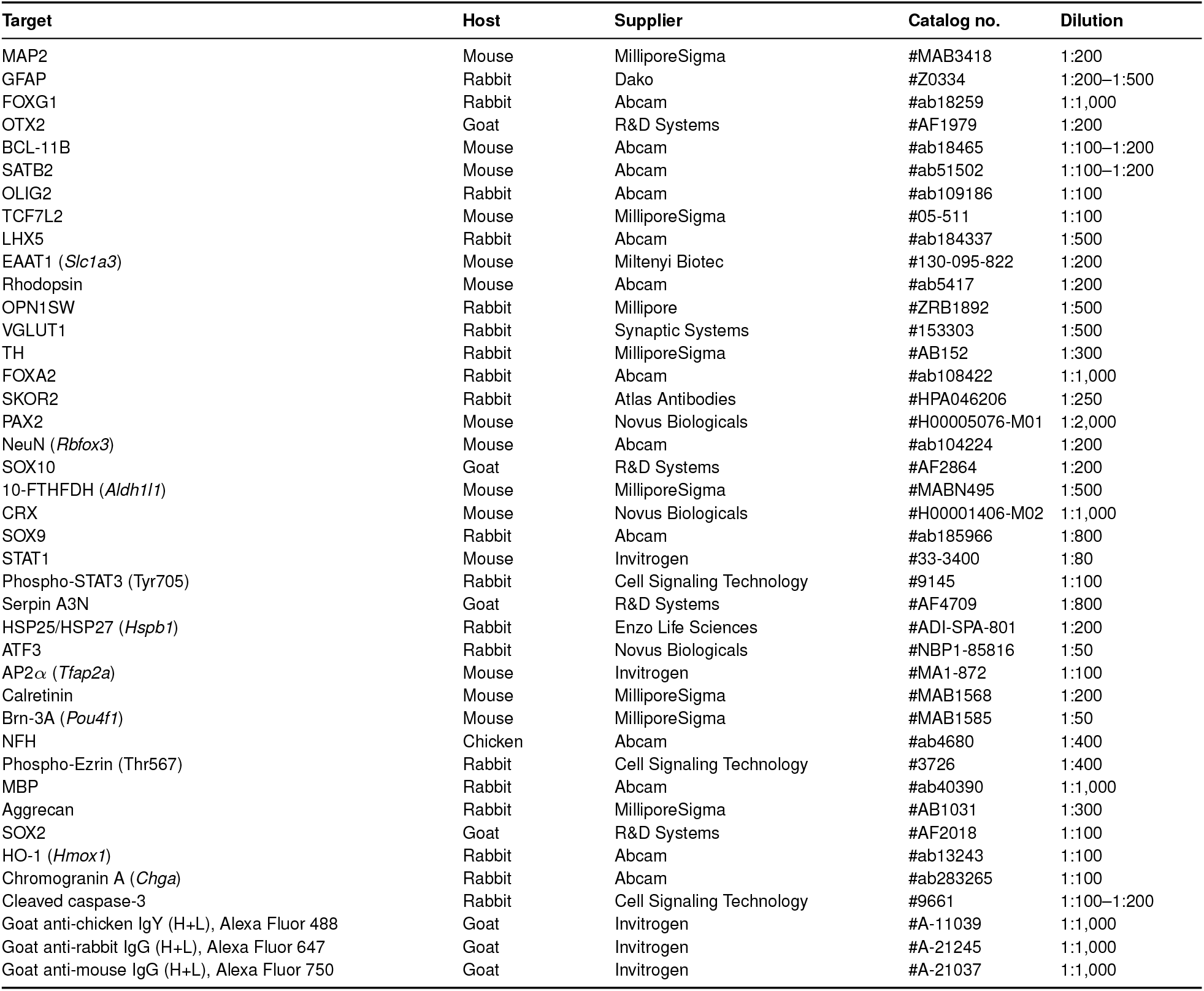

### Immunofluorescence staining, imaging and quantification

Mouse neural organoids were washed in DPBS −/− (Gibco #14190250) and fixed for either 2h at 25°C or 24h at 4°C in 4% Paraformaldehyde (Thermo Scientific #J19943.K2) before the PFA was washed off and the samples were stored in PBS + 0.01% sodium azide (VWR, 786-299) before further processing. Fixed organoids were subsequently embedded, sectioned and imaged as described previously ^123^. In brief, mouse neural organoid slices were embedded in an HistoGel (Epredia, HG-4000 012) mold. The HistoGel block was placed in a biopsy cassette and dehydrated using a fully automated tissue processor (HistoCore PEARL, Leica). The block was then embedded in Paraffin on the Medite Embedding station TES99 and sectioned into 3.5 *µ*m thick sections with a microtome (Thermo Microm HM355S). Multiplexed immunofluorescence staining of the sections was performed using the Ventana Discovery Ultra automated tissue stainer (Roche Tissue Diagnostics) using the settings described previously ^123^. A list of primary antibodies can be found in Supplementary Table 6. Next the corresponding opal dye (Opal 480 FP150001KT, Opal 520 FP1487001KT, Opal 570 FP1488001KT, Opal 620 FP1495001KT, Opal 690 FP1497001KT, Opal 780 FP1501001KT, Akoya Biosciences) were applied, previously prepared following manufacturers instructions. Lastly, samples were counterstained with DAPI (Roche).

For manual stainings the sections were deparaffinized by baking at 60°C for 8 minutes, followed by further heating up to 69°C for 8 minutes for subsequent deparaffinization. This was repeated three times. Heat-induced antigen retrieval was done with Tris-EDTA buffer pH 7.8 at 95°C for a total of 40 min. The sections were then washed several times with ddH_2_O before permeabilization for 5 min with PBST (0.5% Triton-X) at room temperature. After washing, blocking was performed for 1 hour at room temperature in blocking buffer (Dako #X090930-2). The primary antibodies (chicken anti-NFH, rabbit anti-Aggrecan, mouse anti-MAP2) were diluted in blocking buffer (Dako #X090930-2) and incubated at 4°C overnight. After washing the secondary antibody was diluted 1:1000 in blocking buffer and DAPI immunolabeling incubated for 1 hour at room temperature in the dark. We used the following secondary antibodies: goat anti-chicken IgY Alexa Fluor™ 488-conjugated (Invitrogen #A-11039), goat anti-rabbit IgG Alexa Fluor™ 647-conjugated (Invitrogen #A-21245) and goat anti-mouse IgG Alexa Fluor™ 750-conjugated (Invitrogen #A-21037). DAPI (Invitrogen #62248) was used at a 1:500 concentration. Slides were mounted with ProLong Antifade Glass mounting media (Invitrogen #P36982) and the slides were left to cure overnight in the dark.

### Microscopy imaging

The multiplexed, Ventana Discovery Ultra-stained sections were imaged at 20x magnification on the Vectra® Polaris™ (PerkinElmer) automated microscope. Channels unmixing and image tiling was performed using PhenoChart (v1.0.12) and inForm (v2.4). To obtain higher resolution images (Fig. 1e, Fig. 2d) of multiplexed stainings, selected sections were re-imaged at 25x magnification (Leica Microsystems #11506375) using a point-scanning confocal microscope (Leica Stellaris 8). Manually stained sections (Extended Data Fig. 1n) were imaged using a point-scanning confocal microscope (Leica Stellaris 8) at 25x magnification.

### Image analysis

Multiplexed immunofluorescence images were analyzed using HALO software (v3.6.4134.396; Indica Labs). Individual organoids on the sections were isolated using the “Segment TMA” module and annotated according to the treatment condition, with any organoids lacking an epithelium were excluded from the analysis. To classify distinct tissue architectures, a Random Forest classifier was trained using all markers from the multiplexing panel. Retinal organoids were categorized into four classes: “glass” (empty area), “core” (inner part of the organoids with dead cells and non retinal cells), “fold” (folds in the paraffin sections) and “epithelium” (retinal epithelium). Downstream quantification of the retinal organoids was restricted exclusively to data from the class classified as “epithelium”. For brain organoids, the Random Forest algorithm was trained to discriminate between “organoid” and “glass” classes, and downstream analysis was restricted to data from the “organoid” class.

Fluorescent marker abundance was quantified using two HALO Algorithms: For cytoplasmic stainings (EAAT1, RHO, OPN1SW, MAP2, 10-FTHFDH, GFAP, HspB1) the “Area Quantification FL v2.3.4” algorithm was used and the total % of positive area was exported for downstream analysis. For nuclear markers (SOX9, SOX2, SOX10, NeuN, p-STAT3, HO-1, Serpin A3N) quantifications the “CyoNuclear FL v2.0.12” was used and the total % of positive cells were used for downstream quantification. For the nuclear percentages of positive cells (SOX9/p-STAT3, CRX/p-STAT1, CRX/ATF3, DAPI/ATF3) the first nuclear marker was set as “nuclear dye” and the marker to be quantified was set as “Dye 1”. Lookup table (LUT) settings were carefully adjusted on a per-section basis: thresholds were manually set to exclude background noise and maximize visual contrast without over-saturating the target signal. Because of technical variability and per-section LUT adjustments, fluorescence metrics cannot be compared across different sections. To ensure rigorous analysis, all direct comparisons and quantifications shown within a single figure panel derive from the same section; however, intensities should not be compared across different panels. For NFH quantification, a Random Forest classifier was trained to segment continuous NFH^+^ areas (axons) and axon-level shape metrics (area and perimeter) for each organoid was exported and analyzed using R. Axon elongation was computed as Perimeter^2^ / (4*π* * Area) and axons were classified as healthy if their elongation was >4. The number of healthy axons was then normalized by the total axonal area (sum of all axon areas) per organoid. The summary from the HALO analysis was exported and the plots were generated by using ggplot2 (v.4.0.2) in R (v.4.4.2).

### Image visualization

Images from the Vectra Polaris microscope were converted into the OME-Zarr file format and opened in Napari v.0.6.4. LUT were set based on visual assessment and kept constant across direct condition comparisons but not across figures or experiments.

### Metabolomics sample preparation and mass spectrometry analysis

Prior to metabolite extraction, organoids were transferred to a 96-well round-bottom plate to estimate their volume and weight. Bright-field images were acquired using an Olympus CKX53 microscope at 4x resolution, maintaining constant image acquisition settings throughout the time-course. Image processing was automated using a custom Fiji macro to mask the organoids and fit an ellipse. To estimate weight from two-dimensional images, we modeled the organoids as prolate spheroids. Because organoids naturally settle on their flatter side, we applied an empirical correction factor of 0.9 to the minor axis to account for the reduced, unobserved z-axis depth. We further assumed a constant tissue density of 1.08 pg/*µ*m^3^, which corresponds to the average density of brain tissue ^124^. Based on these geometric and physical assumptions, the estimated organoid volume *V* and weight *W* were calculated using the following formulas:

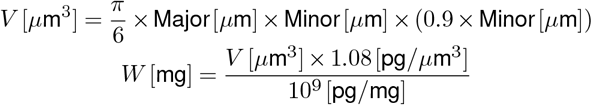

This image-based estimation approach was validated by pooling and physically weighing 28 previously imaged organoids, with the estimated and measured total weights agreeing within a 20% margin. The volume of extraction buffer to be added per sample was then adjusted to normalize sample concentrations to either 25, 50 or 100 *µ*L/mg depending on total input. To rapidly arrest metabolism, samples were quenched under strict temperature control: working one replicate at a time, organoids were transferred using a wide-bore tip into a 1.5 mL tube containing ice-cold DPBS −Ca^2+^/−Mg^2+^ (Gibco #14190250), then immediately moved to tubes containing beating beads (Sarstedt #72.694.406; Cayman Chemical #10402) situated on a −20°C cold block. Residual DPBS −Ca^2+^/−Mg^2+^ (Gibco #14190250) was quickly removed, and an adequate volume of Extraction Buffer (40% methanol, 40% acetonitrile, 20% HPLC-grade H_2_O, supplemented with a 1:100 dilution of 2.5 mM phenylhydrazine) precooled to −20°C was added. Samples were temporarily stored at −20°C during handling and subsequently transferred to −80°C for storage. For final extraction, tissue homogenization was performed using a Precellys homogenizer equipped with a Cryolys cooling unit maintained at 0°C with dry ice. The samples were subjected to three 20-second grinding cycles at 10,000 RPM, separated by 30-second pauses to keep the temperature at 0°C. To isolate soluble polar metabolites from beads, cellular debris, and precipitated proteins, the homogenate was centrifuged at maximum speed for 5 minutes at 4°C. The cleared supernatant was carefully collected, and a 30 *µ*L aliquot was transferred to a 96-well plate for subsequent mass spectrometry analysis.

Samples were analyzed by utilizing flow injection time-of-flight mass spectrometry with an Agilent 6550 iFunnel Q-TOF System as described previously ^125^. 2 *µ*l of sample were injected using a Gerstel MPS Robotic autosampler coupled to an Agilent 1260 Infinity II bio-inert quaternary pump with a constant flow of 0.15 ml/min water/isopropanol mix (40:60, v/v) with 1mM ammonium fluoride, containing two compounds for online mass axis calibration: 3-Amino-1-propanesulfonic acid (M-[H+]-138.0230 m/z, Sigma Aldrich, A76109) and hexakis (1H,1H,3H-tetrafluoropropoxy)phosphazine (M+[F-]-940.0015 m/z, HP-0921, Agilent Technologies, G1969-85001). The ion source settings for ionization in negative mode were kept at 225°C source temperature, flow rate 11 L/min drying gas, 20 psi nebulizer pressure, 350°C sheath gas temperature, 10 L/min sheath gas flow, 3500 V Vcap voltage, 2000 V nozzle voltage, 350 V fragmentor voltage and 750 V octopole voltage. Ms1 profile mass spectra were recorded in a mass range of 50–1000 m/z in 4GHz high-resolution mode with an acquisition rate of 1.5 spectra/s and an acquisition stop time of 0.7 min. Raw ms1 profile mass spectra were converted to open source mz5 format using MSConvert from the Proteowizard tool ^125,126^ and further processed with an in-house data processing environment in MATLAB R2024b (The MathWorks, Natick) to compress data, recalibrate summed profile spectra, adjust baselines and pick centroid ion masses in each sample. Detected ions were putatively annotated by matching mass-to-charge ratios to a reference list of calculated masses of metabolites listed in the Human Metabolome Database (HMDBv5) ^127^ within 0.003 Dalton mass accuracy. The reference mass list was generated from the respective sum formulae, considering deprotonation as the most prevalent mode of ionization in the chosen acquisition conditions. To allow for the annotation of *α*-carboxylic acid derivatives formed in presence of phenylhydrazine in the extraction solvent ^128^, sum formulae for the phenylhydrazone derivatives (+C6H8N2-H2O) of a total of 30 *α*-keto acid compounds (selected via KEGG SimComp search http://www.genome.jp/tools/simcomp/) were added to the metabolite list for annotation. In the absence of prior chromatographic separation, flow injection TOF-MS cannot distinguish isobaric metabolites, as well as in-source fragments that are detected at the identical monoisotopic mass. Downstream computational analysis and quality control were performed in R v4.4.2. Metabolites with more than two missing values were filtered out, and remaining missing values were imputed using a pseudo-count of 1. To eliminate noisy features, metabolites were removed if their average intra-replicate coefficient of variation (CV) exceeded their inter-time-point CV. Principal Component Analysis (PCA) was computed on centered and scaled abundance data. To identify temporal metabolic modules, replicate-averaged abundances were Z-score normalized per metabolite and grouped utilizing k-means clustering (*k* = 3, 25 random starts), with the optimal cluster count confirmed via elbow and silhouette methods. To handle ambiguous feature annotations, mass-to-charge (m/z) values matching multiple putative HMDB IDs were split into separate entries prior to Over-Representation Analysis (ORA). Although this approach artificially inflates the number of input features, it serves as the most comprehensive strategy available to avoid discarding potentially valid biological pathways when handling unresolved isomeric or isobaric metabolites. Over-representation analysis of the resulting metabolite list was then mapped against the HMDB database using the MicrobiomeProfiler package, and a custom Jaccard similarity filter (threshold = 0.95) was applied to prune redundant, overlapping pathways.

### Electrophysiological measurements

Mouse minimally guided brain organoids were cultured as described above and maintained in Differentiation Media with Vitamin A, shaking, with media changes every other day until further processing. On day 13 organoids were washed in ice-cold HBSS with Ca^2+^/Mg^2+^ (Thermofisher, 14025092), embedded in low-gelling temperature agarose 3% (w/v) (Sigma-Aldrich, A9414) at 40°C in peel away embedding molds (Sigma, E6032), and kept on ice for 20 minutes until ready for slicing. Organoids were then sectioned into 300 *µ*m slices, in ice-cold HBSS with Ca^2+^/Mg^2+^, using a Vibratome (Leica VT1000S) with a sterile razor blade, collected onto Millicell-CM cell culture inserts (Merck Millipore, PICM0RG50) in six-well plates and left to equilibrate for 1 hour at 37°C in serum-supplemented slice culture medium (SSSCM): DMEM (Thermo Fisher Scientific, 31966047), 10% FBS, 0.5% (w/v) glucose, supplemented with 0.5% antibiotic-antimycotic (ThermoFisher Scientific, 15240062). SSSCM was then replaced with serum-free slice culture medium (SFSCM): Neurobasal (Thermo Fisher Scientific, 21103049), 2% B-27 supplement with vitamin A (Thermo Fisher Scientific, 17504044), 0.5% (w/v) glucose, 0.5% Glutamax (Thermo Fisher Scientific, 35050061) and antibiotic-antimycotic (ThermoFisher Scientific, 15240062). Slices were cultured at an air-liquid interface (ALI) for at least 48 hours, with daily media changes to improve oxygen and nutrient supply ^116^. High-density microelectrode array (HD-MEA) plates were coated with 0.05% polyethylenimine (PEI, Sigma-Aldrich, 408727) prepared in borate buffer (pH 8.5; Thermo Scientific, 28341) and incubated at 37°C with 5% CO_2_. After 1 hour of incubation, the chips were washed three times with distilled water, left to dry, coated with laminin (0.02 mg/ml; Sigma-Aldrich, 67407300) and incubated overnight at 37°C with 5% CO_2_. The next day organoids were plated and 500 *µ*L of media was carefully exchanged every other day.

Spontaneous electrical activity of developing mouse brain organoids was recorded using high-density microelectrode arrays (HD-MEAs; 6- and 24-well plates, MaxWell Biosystems AG, Zurich, Switzerland). Weekly longitudinal recordings were initiated on day 20 and consisted of measuring the array-wide electrical activity of organoids using the Activity Scan assay (MaxLab, MaxWell Biosystems AG), followed by 30-min network recordings of the intrinsic population activity. We included network recordings from six sliced and six unsliced organoids. All data was spike-sorted using Kilosort 2.5^129,130^ and manually quality-controlled using the phy software suite ^131^. Pairwise spike-train synchrony was quantified for recordings with at least 3 units, by computing cross-correlograms in 1-ms bins over a *±* 20 ms window and summing coincident spikes within that window. The resulting coincidence count was normalized by the geometric mean of the two trains’ spike counts. Axonal conduction velocities (ACVs) were estimated in late-stage organoids (days 35–41) by capturing per-neuron extracellular footprints via the Axon Tracking assay (MaxLab, MaxWell Biosystems AG) using a publicly available axon velocity toolbox ^132^. In brief, the channel-detection threshold was lowered to 1% of the peak amplitudes to ensure recovery of smaller axonal signals. Reconstructed branches were post-processed by splitting any path containing a consecutive peak-latency gap >1 ms and refitting each segment by ordinary least squares against cumulative path distance. We discarded segments with fewer than 5 electrodes or *R*^2^ *<* 0.9. Within each well, cross-neuron duplicate branches arising from over-segmentation by upstream footprint extraction were merged using a 50-*µ*m spatial-bin Jaccard index *≥* 0.5 (highest-*R*^2^ branch retained); only branches from footprints with completeness *≥* 0.5 were included in pooled statistics. Pharmacological blockade of ionotropic glutamate receptors was performed in late-stage brain organoids (day 50, *n* = 4) to confirm the presence of actively coupled, functionally mature excitatory synapses. Spontaneous population activity was recorded for a 30-minute baseline period, followed by a subsequent 30-minute recording during the acute application of the competitive NMDA receptor antagonist AP5 (20 *µ*M) and the AMPA/kainate receptor antagonist CNQX (20 *µ*M).

### Dose selection by qualitative cytotoxicity assessment

The dose used for every perturbation was carefully tested. In short, an initial dose was selected based on previously reported use cases and concentrations ranging from 3 to 100 fold higher and lower were tested alongside to find the dose at which a molecule starts to show toxic effects. For all H_2_O_2_ treatments we first removed known antioxidizing agents (L-Ascorbic Acid, Vitamin A and 2-mercaptoethanol) from maintenance media. If not otherwise mentioned, toxicity was assessed by qualitative visual assessment of bright-field images, using criteria such as overall morphology, change in tissue density (e.g. loss of bright epithelium), and cell shedding. All dose testing experiments were performed on day 25–28 retinal organoids and day 30 brain organoids. All tested doses, toxicity assessments and final selected doses are available in Supplementary Table 3. For the OSM pathway inhibition experiments, the selected doses after bright-field assessment and cleaved-Caspase 3 staining were 0.5 *µ*M for TPCA-1 (MedChemExpress, #HY-10074) and 10 *µ*M for Ruxolitinib (MedChemExpress, #HY-50856).

### Statistical analysis

All statistical analyses were performed using R version 4.4.2. Tests for differences in average proportions across groups were conducted using Welch’s t-tests after applying an arcsine transformation to the proportions. Cohen’s h was computed as the difference between the arcsine-transformed means of each group to estimate the effect size for these comparisons. For comparisons of signature scores, we computed Cohen’s d to estimate effect size. Unless otherwise mentioned, all other comparisons in this study utilized unpaired Welch’s t-tests. To account for multiple testing, p-values were adjusted using the Benjamini-Hochberg false discovery rate (FDR) method, with an FDR < 0.05 considered statistically significant. All box plots display median and interquartile range.

## Data and code availability

All sequencing, image analysis, metabolomic, electrophysiological data and code used in this study will be made publicly available via GitHub and Mendeley Data upon publication. During the peer-review process, these materials can be provided confidentially upon request. Please contact the corresponding authors for inquiries.

## Supplementary Information

**Extended Data Figure 1.**
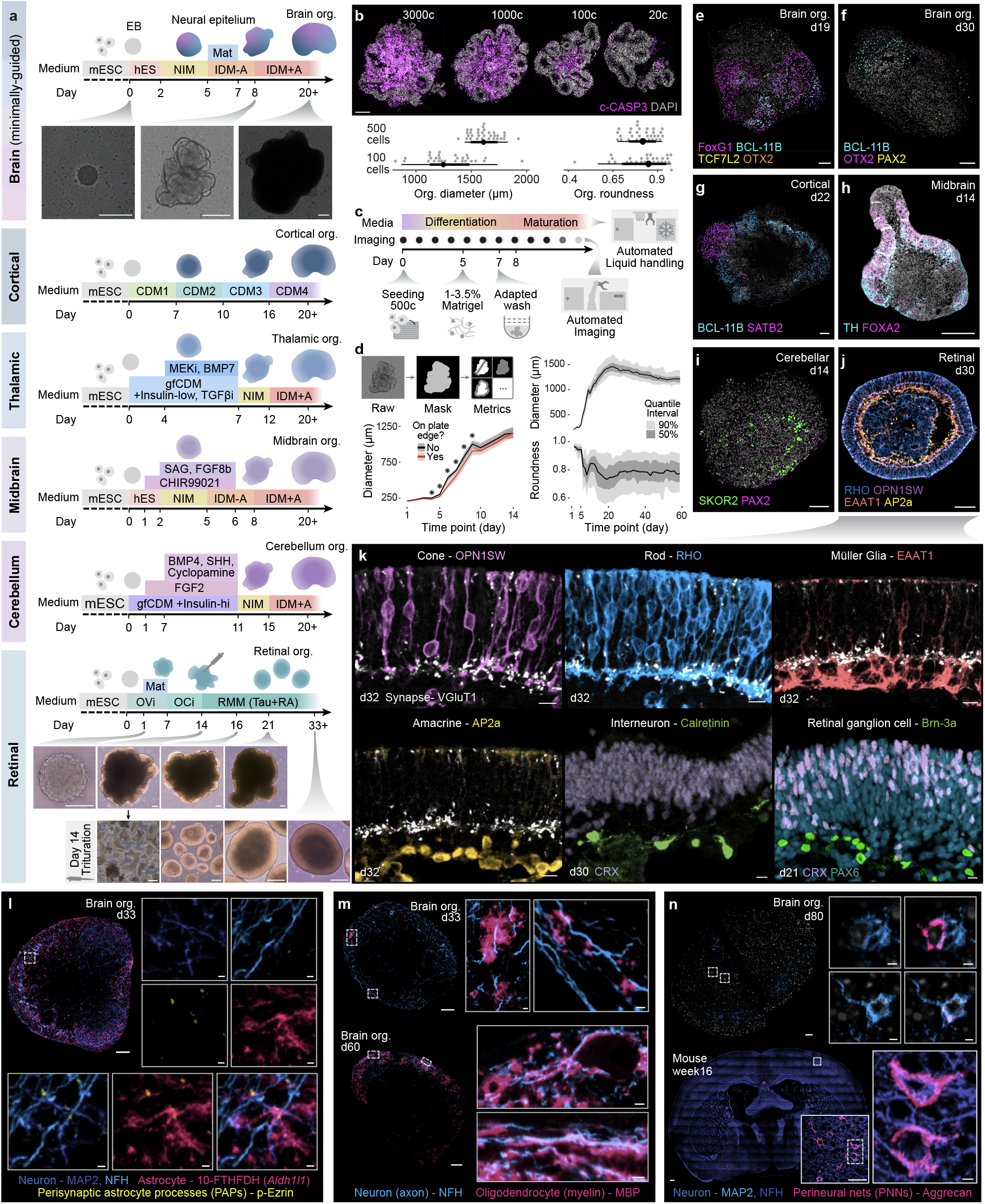
Mouse neural organoids growth, cellular composition and neuro-macroglial interactions. **(a)** Overview of mouse neural organoid protocols. Insets display representative bright-field images. Scale bars, 200 *µ*m. **(b)** Optimization of cell-seeding density. Top: Representative immunostainings of apoptosis marker cleaved-Caspase3 in day 7 mouse brain organoids initiated from 3000, 1000, 100 and 20 cells. Bottom: distribution of diameter and roundness for day 20 organoids initiated from 500 and 100 cells. Organoids started from 100 cells showed greater diameter variability (SD = 244 *µ*m) compared to those from 500 cells (SD = 127 *µ*m) (bottom). Median, 66% and 95% quantile intervals displayed. Scale bar, 100 *µ*m. **(c)** Overview of equipment and protocol modifications for automation. Changes included starting from 500 cells, providing Matrigel exposure as 1–3.5% Matrigel in solution, and using *≥* 3 consecutive half media changes with cold medium on day 7. **(d)** Extraction of morphological features from automated brightfield imaging to track organoid development. Top left: Pipeline overview. Right: Rapid growth and decreased roundness occur post-Matrigel addition (day 5) and peaks near day 20. Bottom left: Growth curves stratified by plate position display a transiently reduced diameter in edge-well organoids between days 4–9. Eight-spoked asterisks (✳) denote significant and large effects (p < 0.01; Cohen’s d > 1. **(e-j)** Immunostainings of regional markers in mouse neural organoids guided towards the following regional identities: **(e, f)** brain (pan-forebrain FoxG1; cortical BCL-11B; dien-mesencephalic TCF7L2, OTX2; hindbrain PAX2); **(g)** cortical (deep/upper layer BCL-11B, SATB2); **(h)** midbrain (dopaminergic TH, floor plate FOXA2); **(i)** cerebellar (Purkinje SKOR2, hindbrain PAX2); and **(j)** retinal (photoreceptor RHO/OPN1SW, amacrine AP2a). **(j, k)** Representative immunostainings of major cell types forming a stratified epithelium in mouse retinal organoids. **(j)** Whole-organoid cross-section of a day 30 organoid co-stained for photoreceptor (OPN1SW; Rhodopsin), glial (EAAT1) and amacrine (AP2a) markers. Scale bars: 100 *µ*m. **(k)** High-magnification views displaying cones (OPN1SW), rods (Rhodopsin), Müller glia (EAAT1), amacrine cells (AP2a) and glutamatergic synapses (VGluT1) in day 32 organoids; interneurons (Calretinin) and photoreceptors (CRX) in day 30 organoids; and retinal ganglion cells (Brn-3a) and progenitors (PAX6) in day 21 organoids. Scale bars: 10 *µ*m. **(l-n)** Advanced cellular maturation and interactions in brain organoids. **(l)** Tripartite synapse interactions between neurons (MAP2, NFH) and perisynaptic astrocyte processes (p-Ezrin, 10-FTHFDH) in day 33 brain organoid. Scale bars: 100 *µ*m; insets 5 *µ*m. **(m)** Myelinating oligodendrocytes (MBP) wrapping neuronal axons (NFH) in day 33 and 60 brain organoids. Scale bars: 100 *µ*m; insets 5 *µ*m. **(n)** Neurons enveloped by perineuronal nets (Aggrecan) in day 80 brain organoid (top), mirroring primary mouse brain structures at week 16 (bottom). For brain organoids, left insets show only MAP2 signal and right insets show overlaid MAP2 and Aggrecan signals. Scale bars: 50 *µ*m; insets 5 *µ*m. mESC = mouse embryonic stem cell, EB = Embryoid Bodies, org. = organoid, Mat = Matrigel.

**Extended Data Figure 2.**
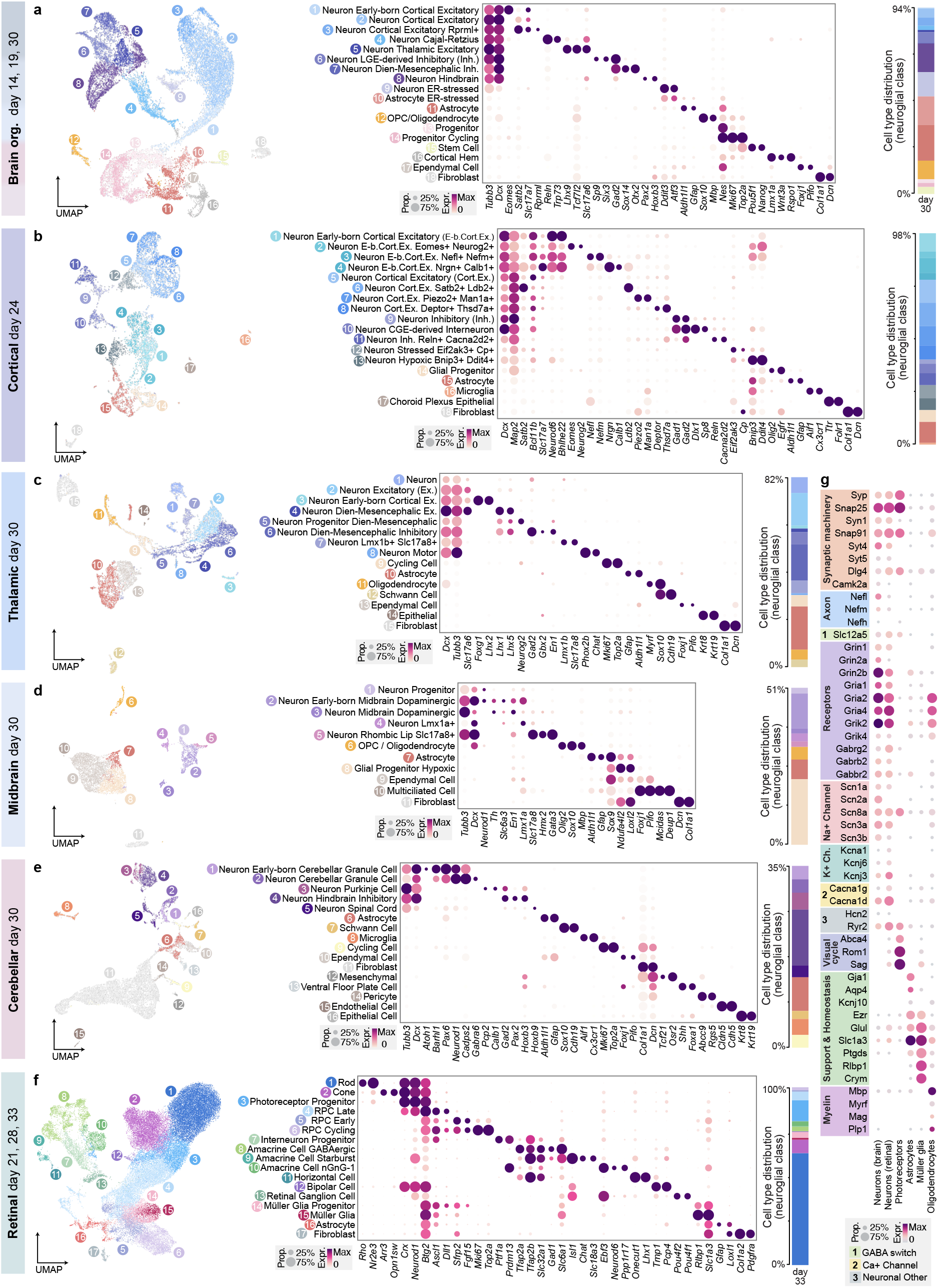
Transcriptomic profiling of mouse neural organoids. **(a-f)** Transcriptomic characterization of mouse neural organoids patterned toward distinct CNS regions. Left: UMAP embeddings colored by annotated cell type. Middle: Dot plots of cell-type-specific marker expression supporting annotation (dot color indicates scaled average expression; dot size indicates the proportion of expressing cells). Right: Stacked bar plots displaying relative proportions of neuroglial cell types. **(a)** Integrated brain organoid datasets from day 14, 19 and 30, containing cortical excitatory neurons (*Eomes, Satb2, Slc17a7*), alongside Cajal-Retzius cells (*Reln, Trp73*), thalamic excitatory neurons (*Lhx9, Tcf7l2, Slc17a6*), LGE-derived inhibitory neurons (*Sp9, Six3, Gad2*), diencephalic-mesencephalic inhibitory neurons (*Sox14, Otx2*), and hindbrain neurons (*Pax2, Hoxb3*). **(b)** Cortical organoids containing cortical excitatory neurons (*Satb2, Bcl11b, Slc17a7*) and inhibitory neurons, notably CGE-derived interneurons (*Dlx1, Sp8*). **(c)** Thalamic organoids comprising target diencephalic neurons (*Lhx1, Lhx5*) alongside developmentally adjacent telencephalic (*Foxg1, Lhx2*) and mesencephalic (*En1*) subtypes. **(d)** Midbrain organoids containing midbrain dopaminergic (*Th, Slc6a3*) and rhombic lip glutamatergic (*Slc17a8*) neurons, but yielding 49% of off-target cell types. **(e)** Cerebellar organoids comprising cerebellar granule (*Atoh1, Barhl1, Pax6*) and Purkinje (*Pcp2, Calb1*) cells, alongside hindbrain/spinal cord neurons (*Pax2, Hoxb9*), but yielding 65% of off-target cell types. **(f)** Integrated retinal organoids datasets (day 21, 28 and 33) containing diverse retinal cell types, including photoreceptors (*Rho, Arr3*), amacrine (*Tfap2a*), horizontal (*Onecut1*), retinal ganglion (*Pou4f1*) and bipolar cells (*Trnp1, Pcp4*), alongside Müller glia (*Rlbp1, Slc1a3*). **(g)** Dot plot displaying expression of maturation markers across target cell classes at advanced time points (dot color indicates scaled average expression; dot size indicates the proportion of expressing cells).

**Extended Data Figure 3.**
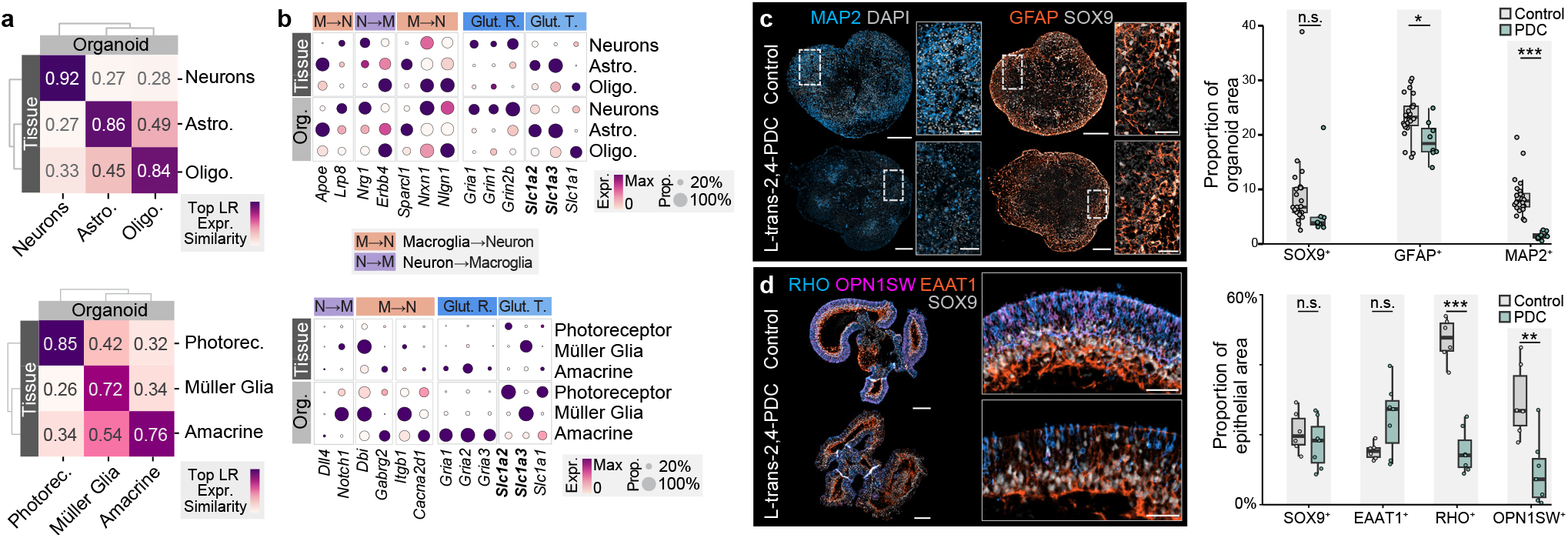
Neuro-macroglial interactions in mouse neural organoids. **(a, b)** Transcriptome-based assessment of neuro-macroglial interactions. **(a)** Heatmaps displaying cosine similarity of top receptor-ligand (R-L) pair expression between primary tissue and organoids for brain (top) and retina (bottom). **(b)** Dot plots validating the shared compartmentalized expression of classic neuroglial interacting pairs (e.g., *Apoe → Lrp8, Dll4 → Notch1*) between organoids and primary reference for brain (top) and retina (bottom). **(c, d)** Inhibition of glutamate transporter with the broad-spectrum EAAT inhibitor L-trans-2,4-PDC (PDC). **(c)** Representative immunostainings (left) and quantification (right) of day 33 PDC-treated brain organoids. PDC exposure results in a significantly decreased neuronal MAP2^+^ area (Cohen’s h = −0.35) and astrocytic GFAP^+^ area (Cohen’s h = −0.11) but no significant change in astrocytic SOX9^+^ (Cohen’s h = −0.12) area. **(d)** Representative immunostainings (left) and quantification (right) of day 27 retinal organoids treated with PDC for 48h. PDC treatment leads to a reduction in photoreceptor RHO^+^ (Cohen’s h = −0.72) and OPN1SW^+^ area (Cohen’s h = −0.60), whereas Müller glia SOX9^+^ or EAAT1^+^ area are not significantly changed. Points represent independent organoid replicates; *\*\*\** p < 0.001, *\*\** p < 0.01, *\*\** p < 0.05. Photorec. = Photoreceptors; Glut. R. = Glutamate Receptor; Glut. Tr. = Glutamate Transporter; PDC = L-trans-2,4-PDC; n.s. = non-significant.

**Extended Data Figure 4.**
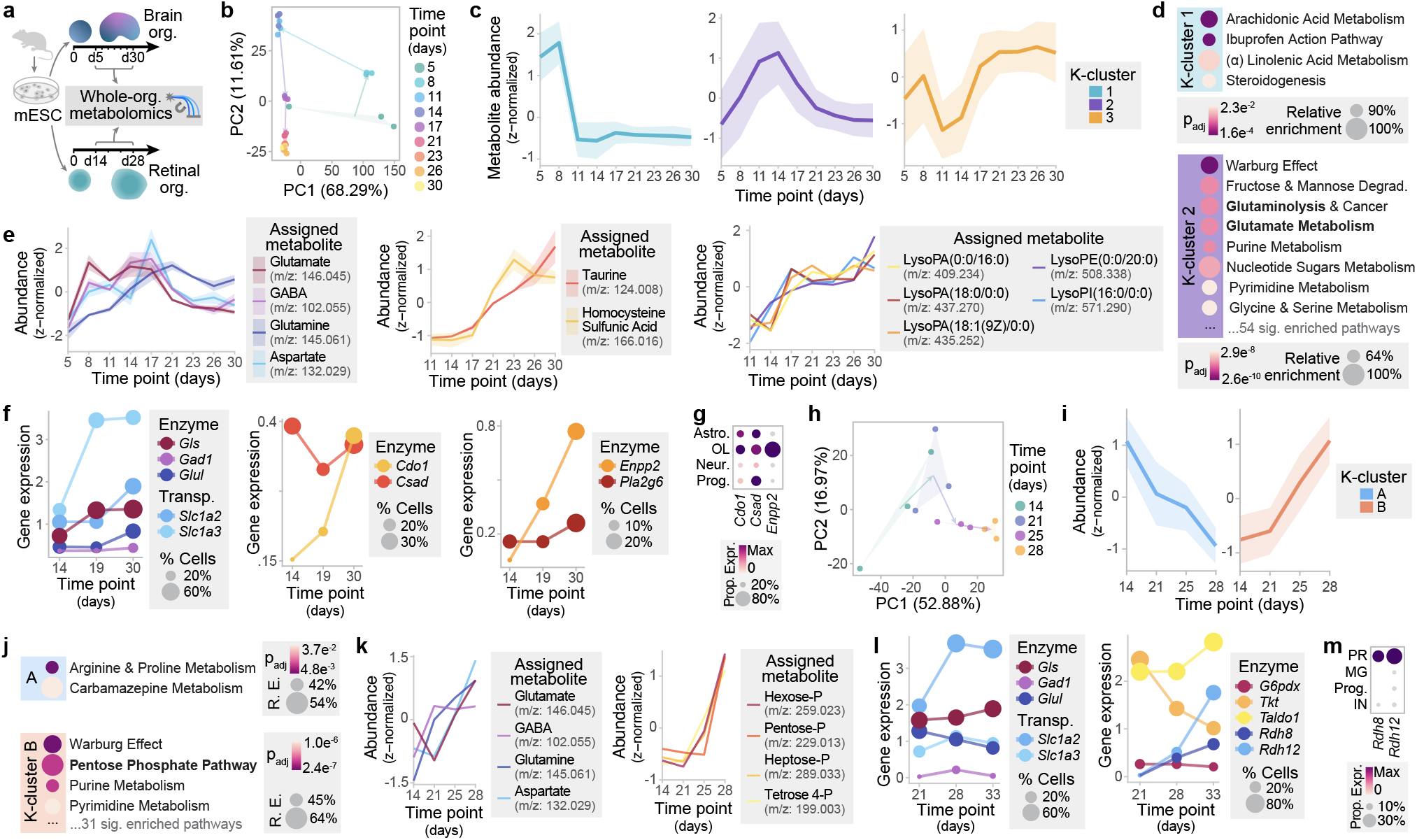
Dynamic metabolic shifts accompanying the maturation of mouse neural organoids. **(a)** Overview of experimental design. Whole-organoid metabolomics of mouse brain and retinal organoids was performed across dense developmental time courses (brain: days 5, 8, 11, 14, 17, 21, 23, 26, 30; retinal: days 14, 21, 25, 28). **(b)** Principal Component Analysis (PCA) of brain organoid metabolomic profiles colored by time point. Arrows reflect transitions between successive time point centroids. Points represent independent biological replicates. **(c)** K-means clustering of brain organoid metabolites by temporal abundance profiles, revealing distinct temporal profiles characterized by early, intermediate, and late relative abundance. Plots display the z-score-normalized average abundance of metabolites across the three identified k-clusters (mean + SD shown). **(d)** Metabolic pathway over-representation analysis for brain organoid k-clusters. Dot size reflects relative enrichment (percentage of pathway metabolites in k-cluster relative to total number of metabolites from this pathway detected); color indicates adjusted p-value. No pathway was significantly over-represented for k-cluster 3. **(e)** Z-score-normalized temporal abundance of selected metabolites across brain organoid development. Left: intermediate-stage accumulation of neurotransmitters and their precursors (glutamate, GABA, glutamine, aspartate). Middle: late-stage enrichment of taurine and its precursor homocysteine sulfinic acid. Right: late-stage enrichment of lysophospholipids. **(f)** Temporal gene expression (pseudo-bulk, log-normalized) and expressing cell fraction (dot size) of metabolic enzymes in brain organoids supporting the metabolomic shifts. Left: neurotransmitter metabolism (*Gls, Gad1, Glul*) and glutamate transporters (*Slc1a2, Slc1a3*). Middle: taurine biosynthesis (*Cdo1, Csad*). Right: lysophospholipid processing (*Enpp2, Pla2g6*). **(g)** Dot plot detailing the cell-type-specific expression of metabolic enzymes (*Cdo1, Csad, Enpp2*) in day 19–30 brain organoids, highlighting compartmentalized expression in astrocytes (Astro.) and oligodendrocytes (OL). **(h)** PCA of retinal organoid metabolomic profiles colored by time point. Arrows reflect transitions between successive time point centroids. Points represent independent biological replicates. **(i)** K-means clustering of retinal organoid metabolites by temporal z-score-normalized abundance profiles (mean + SD shown). **(j)** Pathway over-representation analysis for retinal organoid k-clusters A and B. Dot size reflects relative enrichment (percentage of pathway metabolites in k-cluster relative to total number of metabolites from this pathway detected); color indicates adjusted p-value. **(k)** Z-score-normalized temporal abundance of selected metabolites across brain organoid development. Left: accumulation of neurotransmitters and their precursors (glutamate, GABA, glutamine, aspartate). Right: late-stage accumulation of pentose phosphate pathway (PPP) intermediates. **(l)** Temporal gene expression (pseudo-bulk, log-normalized) and expressing cell fraction (dot size) of metabolic enzymes in retinal organoids supporting the metabolomic shifts. Left: neurotransmitter metabolism enzymes (*Gls, Gad1, Glul*) and glutamate transporters (*Slc1a2, Slc1a3*). Right: PPP enzymes (*G6pdx, Tkt, Taldo1*) and the NADPH-dependent visual cycle enzymes *Rdh8* and *Rdh12*. **(m)** Dot plot detailing the cell-type-specific expression of NADPH-dependent enzymes in day 28–33 retinal organoids, highlighting compartmentalized expression in photoreceptors (*Rdh8, Rdh12*). mESC = mouse embryonic stem cell; Astro. = Astrocyte; OL = Oligodendrocyte; Neur. = Neuron; Prog. = Progenitor; R. E. = Relative enrichment; PR = Photoreceptor; MG = Müller glia; IN = Interneuron.

**Extended Data Figure 5.**
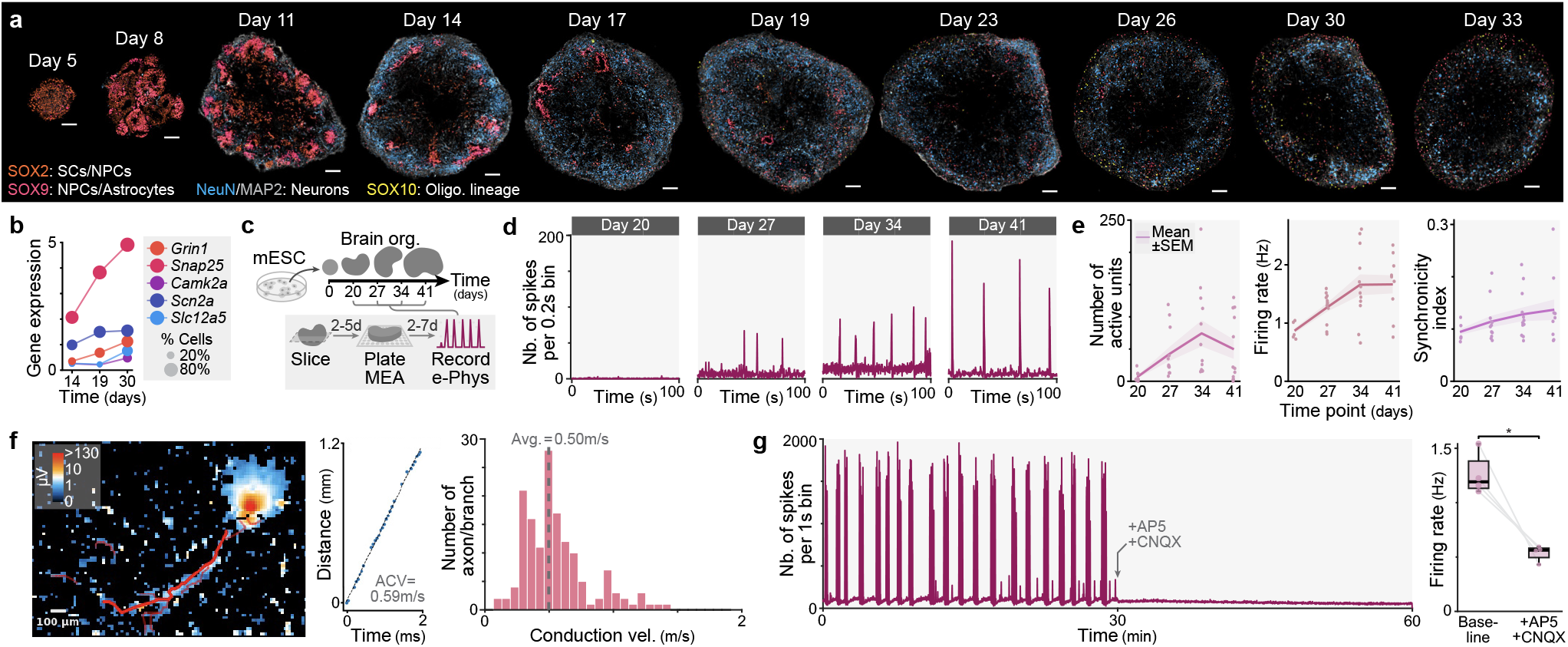
Electrophysiological characterization of mouse neural organoid maturation. Representative immunostainings of neural progenitors (SOX2), neurons (NeuN, MAP2), and oligodendrocytes (SOX10) in brain organoids from day 5 to day 33. SOX9 marks both neural progenitors (organized in rosettes) and astrocytic populations (sparsely distributed). Scale bars: 100 *µ*m. (**b**) Temporal gene expression (log-normalized, pseudo-bulk over neuron subset) and expressing cell fraction (dot size) of electrophysiological genes in brain organoid neurons: excitatory glutamatergic receptor (*Grin1*), synaptic vesicle machinery (*Snap25*), voltage-gated sodium channel (*Scn2a*), mature postsynaptic marker (*Camk2a*), and the chloride transporter driving the GABA switch (*Slc12a5*). **(c)** Experimental workflow for longitudinal electrophysiology. Developing brain organoids were plated onto high-density microelectrode arrays (HD-MEAs), and network activity was recorded between day 20 and day 41. **(d)** Raster plots of spontaneous electrical activity (spikes per 0.2s bin) recorded over a 100-second window across four developmental time points for a representative organoid. **(e)** Quantification of network maturation metrics (number of active units, average neuronal firing rate, and synchronicity index) from HD-MEA recordings (*n* = 12 organoids; 6 intact and 6 sliced). Points represent independent organoid replicates. **(f)** Extracellular mapping of axonal conduction velocity in late-stage organoids (days 35–41). Left: Representative extracellular electrical footprint of a single neuron tracking action potential propagation along an axonal branch (red line). Middle: Linear relationship between cumulative path distance and peak latency for a representative branch, yielding an axonal conduction velocity (ACV) of 0.59 m/s. Right: Histogram of ACVs across 152 reconstructed axonal branches (from 100 successfully tracked neurons across 18 organoids), revealing a median conduction velocity of 0.50 m/s (median *R*^2^ = 0.98; median path length 355 *µ*m). **(g)** Pharmacological validation of glutamatergic network activity. Left: Representative raster plot from late-stage organoid (day 50) before and during the application of the AMPA/kainate and NMDA receptor antagonists CNQX and AP5. Right: Quantification showing a significant reduction in overall spontaneous firing rates (−53%) following CNQX/AP5 application (*n* = 4 organoids). Points represent independent organoid replicates; *\*\** p < 0.05 (paired Student’s t-test). HD-MEA = High-density microelectrode array; SCs = Stem cells; NPCs = Neural progenitor cells; ACV = Axonal conduction velocity; CNQX = 6-cyano-7-nitroquinoxaline-2,3-dione; AP5 = D-2-amino-5-phosphonovalerate.

**Extended Data Figure 6.**
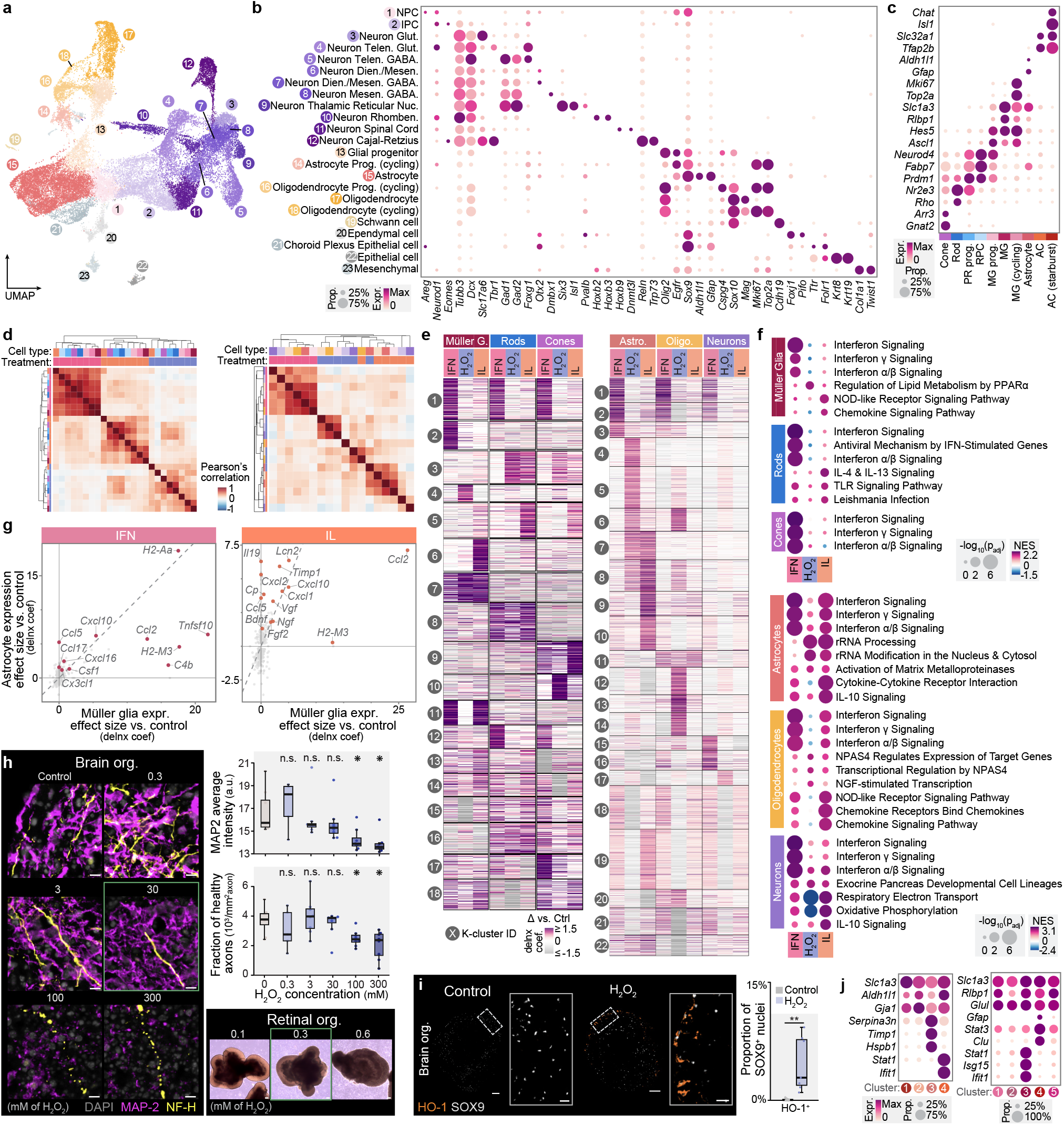
Characterization of perturbed mouse neural organoids and stressor-induced transcriptomic programs. UMAP embedding of brain organoid datasets colored by cell (sub)type. **(b, c)** Dot plots of cell-type-specific marker expression supporting annotation for brain **(b)** and retinal organoids **(c)**. Dot color indicates scaled average expression; dot size reflects the proportion of expressing cells. **(d)** Hierarchically clustered heatmap displaying correlation (Pearson’s r) in treatment-induced gene expression changes across cell types and perturbations for retinal (left) and brain organoids (right). **(e)** K-means-clustered heatmap displaying degree of change in expression (delnx effect size estimates) for all genes differentially expressed upon IFN, H_2_O_2_, and IL treatments across major retinal (left) and brain (right) cell types. **(f)** Dot plot displaying top gene sets significantly upregulated across treatments and main cell types in retinal (top) and brain organoids (bottom) from Gene Set Enrichment Analysis (GSEA). Dot color indicates Normalized Enrichment Score (NES); dot size reflects log-transformed significance; the top 3 gene sets by NES are displayed for each treatment and cell type. **(g)** Scatter plot relating degree of change in expression (delnx effect size estimates) of ligand genes in retinal Müller glia and brain astrocytes relative to control across IFN and IL treatment. **(h)** Dose testing for hydrogen peroxide (H_2_O_2_). Left and top right: Representative immunostainings (left) and quantification (top right) of neuronal cytoskeleton marker MAP2 and axonal marker NFH in H_2_O_2_-treated brain organoids. Points represent independent organoid replicates; asterisks (*\**) denote significant and large effects (p < 0.05; Cohen’s d > 1); scale bar, 10 *µ*m. Bottom right: Representative bright-field images of H_2_O_2_-treated retinal organoids, highlighting epithelial rim darkening and increased density of dead cells in media at higher H_2_O_2_ concentrations. **(i)** Representative immunostainings (left) and quantification (right) of oxidative stress marker HO-1 in control and H_2_O_2_-treated brain organoids. Astrocytes (SOX9^+^) exhibit significant upregulation HO-1. Points represent independent organoid replicates; *\*\** p < 0.01; Cohen’s h = −0.36; scale bars: 100 *µ*m; insets, 20 *µ*m. **(j)** Dot plots of marker gene expression supporting Müller glia and astrocytes subcluster annotation in Fig. 3h-k. Dot color indicates scaled average expression; dot size reflects the proportion of expressing cells.

**Extended Data Figure 7.**
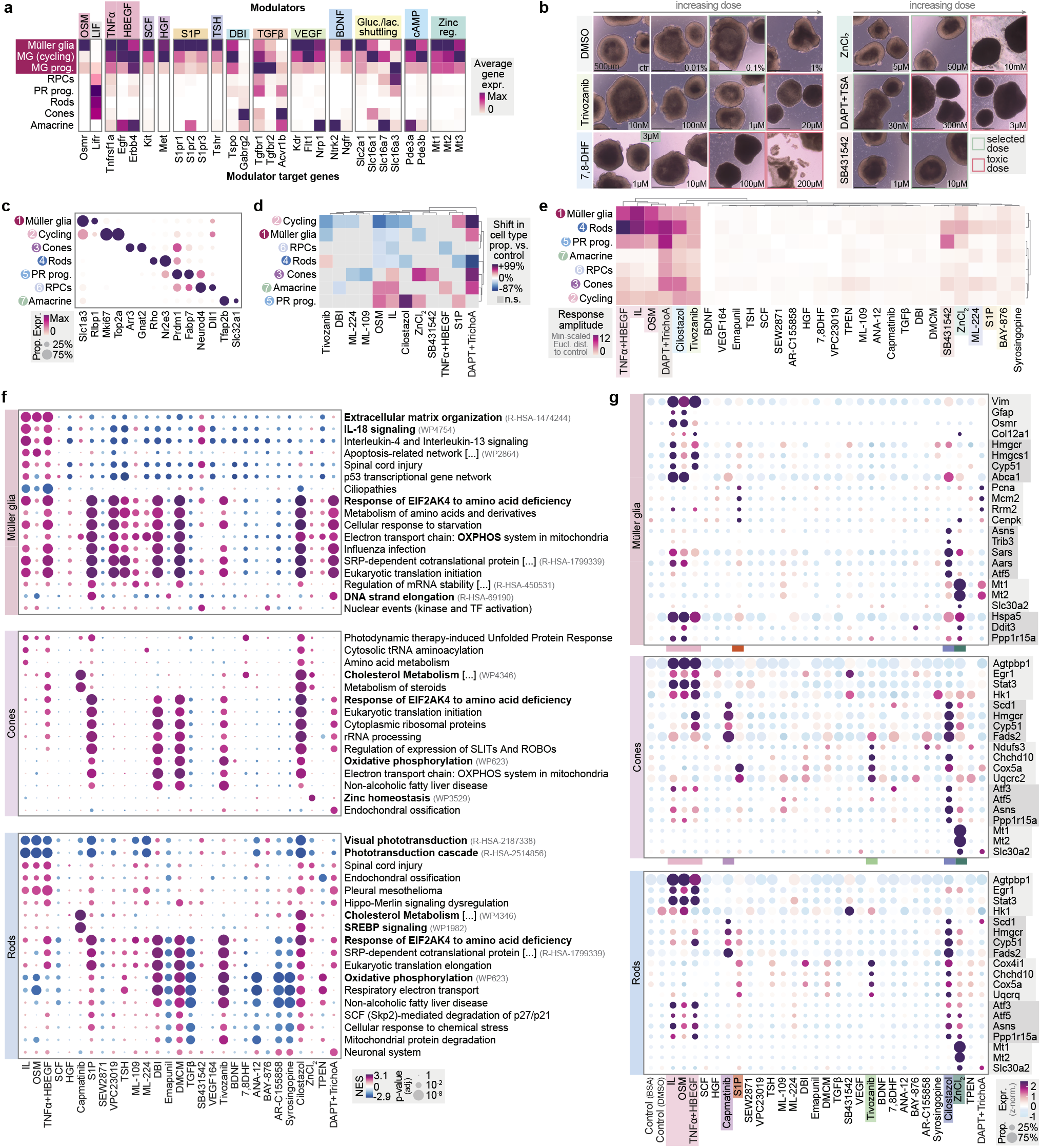
Extended profiling of the Müller glia-targeted single-cell perturbation atlas. **(a)**Heatmap depicting average gene expression in control retinal organoids across cell types of the molecular targets of the selected pathway modulators. **(b)**Representative bright-field images from systematic dose-optimization (examples shown for 6 selected treatments including control). Cytotoxicity was qualitatively assessed over 48h based on organoid morphology, change in tissue density, and dead cell shedding. **(c)** Dot plot of cell-type-specific marker expression supporting annotation. Dot color indicates zero-max-scaled average expression; dot size reflects the proportion of expressing cells. **(d)** Hierarchically clustered heatmap detailing the fold-change (FC) in cell type proportions relative to vehicle controls for perturbations yielding a significant shift (p > 0.05) in ≥ 1 cell type. **(e)** Hierarchically clustered heatmap detailing transcriptomic response amplitude (min-scaled Euclidean distance to control) across all cell types and perturbations. **(f)** Dot plot displaying top significant and treatment(s)-specific gene sets upregulated across Müller glia, cones, and rods from Gene Set Enrichment Analysis (GSEA). Dot color indicates Normalized Enrichment Score (NES); dot size reflects log-transformed significance. **(g)** Dot plot of enriched gene set marker expression across perturbations, including markers for: glial reactivity and ECM remodeling markers (*Vim, Gfap, Osmr, Col12a1*); sterol regulation (*Hmgcr, Hmgcs1, Cyp51, Abca1, Scd1, Fads2*); DNA replication (*Pcna, Mcm2, Rrm2, Cenpk*); integrated stress response (*Asns, Trib3, Sars, Aars, Atf5, Hspa5, Ddit3, Ppp1r15a, Atf3*); zinc homeostasis (*Mt1, Mt2, Slc30a2*); OXPHOS (*Cox4i1, Ndufs3, Cox5a*) and photoreceptor degeneration (*Agtpbp1, Egr1, Stat3,Hk1*). Dot color indicates z-scaled average expression; dot size reflects the proportion of expressing cells. MG = Müller glia; RPCs = Retinal progenitor cells; PR prog = Photoreceptor progenitors; Gluc./lac. = Glucose/lactate; cAMP = cyclic AMP; reg. = Regulation; expr. = expression; NES = Normalized Enrichment Score; Eucl. dist. = Euclidean distance; ISR = Integrated Stress Response; OXPHOS = Oxidative phosphorylation; SREBP = Sterol Regulatory Element-Binding Protein; FC = Fold change; IL = Inflammatory cytokines; OSM = Oncostatin M; TrichoA = Trichostatin A.

**Extended Data Figure 8.**
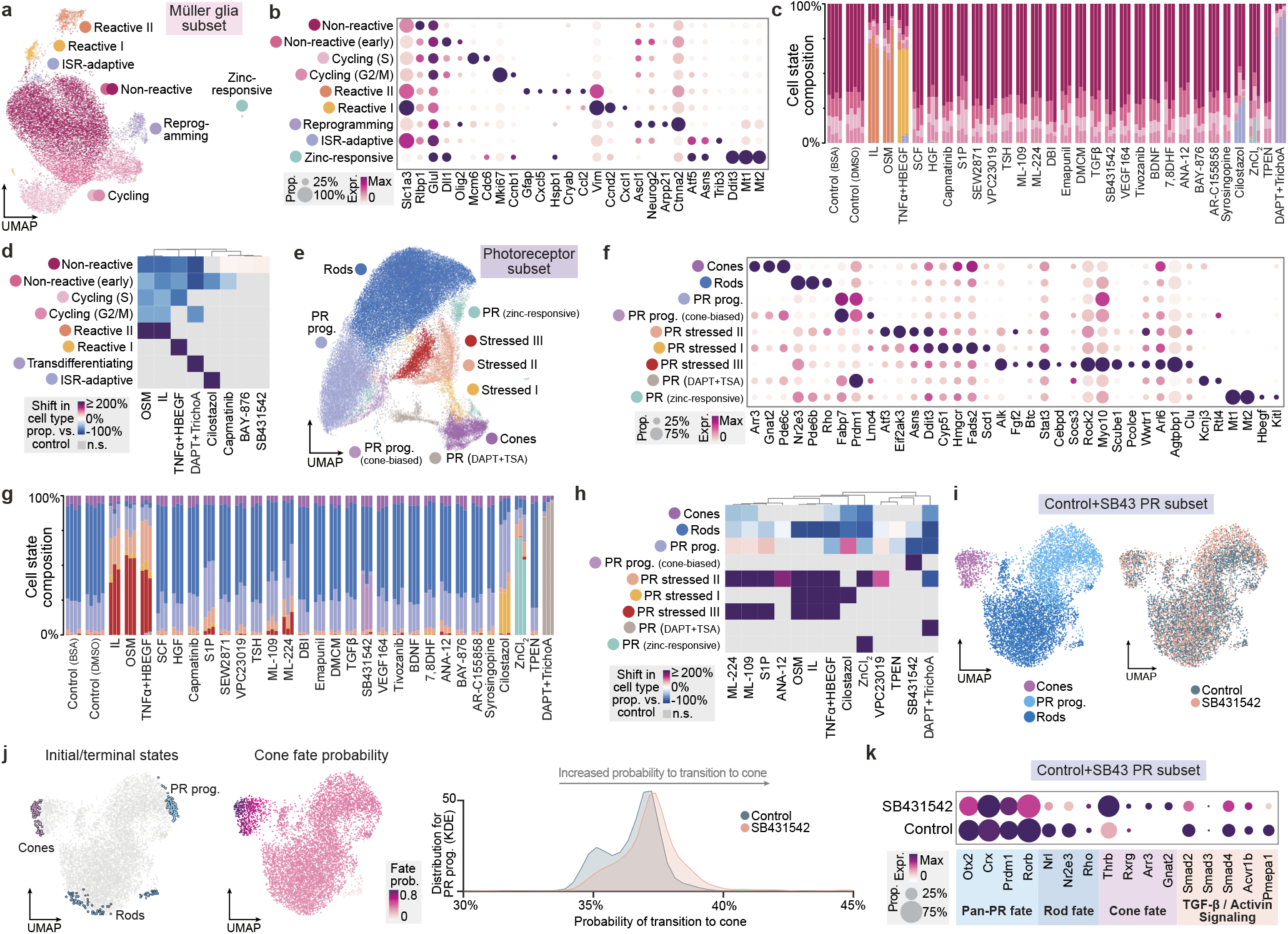
Glial and photoreceptor states in perturbation atlas. **(a)** UMAP embedding of the integrated perturbation atlas subsetted to Müller glia, colored by cell states. **(b)** Dot plot of cell-state-specific marker expression supporting Müller glia state annotation. Dot color indicates scaled average expression; dot size reflects the proportion of expressing cells. **(c)** Stacked bar chart of Müller glia state composition across all conditions; each bar represents an independent biological replicate. **(d)** Hierarchically clustered heatmap detailing the fold-change (FC) in Müller glia state proportions relative to vehicle controls for perturbations yielding a significant shift (p > 0.05) in *≥* 1 state. **(e)** UMAP embedding of the integrated perturbation atlas subsetted to photoreceptors (photoreceptor progenitors, rods and cones), colored by cell states. **(f)** Dot plot of cell-state-specific marker expression supporting photoreceptor state annotation. Dot color indicates scaled average expression; dot size reflects the proportion of expressing cells. **(g)** Stacked bar chart of photoreceptor state composition across all conditions; each bar represents an independent biological replicate. **(h)** Hierarchically clustered heatmap detailing the fold-change (FC) in photoreceptor state proportions relative to vehicle controls for perturbations yielding a significant shift (p > 0.05) in *≥* 1 state. **(i)** UMAP embeddings of photoreceptors from control and SB431542 condition colored by cell type (left) and condition (right). **(j)** UMAP embeddings highlighting calculated marcostates (left) and cone fate probability (middle). Kernel density estimation plot (right) of cone fate probability from photoreceptor progenitor states for SB431542 and control cells. The x-axis is truncated at 0.45 to better visualize the main density peaks; maximum calculated transition probabilities extended to 0.6 for a negligible fraction of outliers. **(k)** Dot plot of rod-cone fate markers and TGF-*β* signaling genes expression. Dot color indicates scaled average expression; dot size reflects the proportion of expressing cells.

**Extended Data Figure 9.**
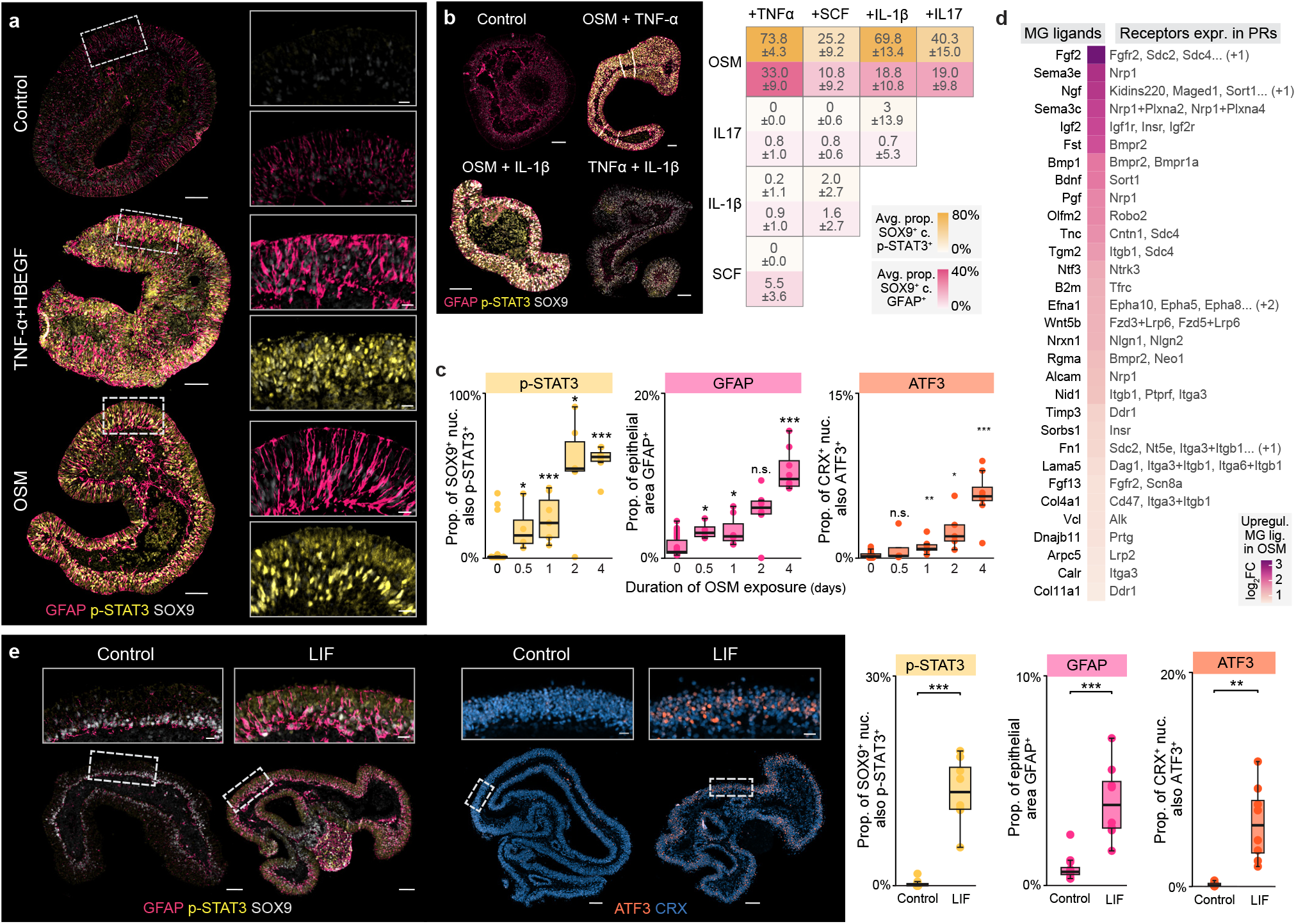
Extended profiling of cytokine-induced retinal phenotypes. **(a)**Representative immunostainings of mouse retinal organoids treated with OSM or TNF*α*+HBEGF for 48h, showing increased levels of p-STAT3 and GFAP in Müller glia (SOX9^+^) nuclei. Corresponding quantification is provided in Fig. 5c. Scale bars: 100 *µ*m; insets, 20 *µ*m. **(b)** Representative immunostainings (left) and quantification (right) of glial reactivity markers p-STAT3 and GFAP in retinal organoids treated with combinatorial pairs of select inflammatory cytokines (Supplementary Table 4) for 48h. Displayed values represent the average *±* standard deviation across 6–8 organoid replicates. Scale bars: 100 *µ*m. **(c)** Immunostaining quantification over 4-day exposure to OSM. Points represent independent organoid replicates; asterisks denote significance compared to day 0 (*\*\*\** p < 0.001, *\*\** p < 0.01, *\*\** p < 0.05). The data displayed in this panel and in Fig. 5h were acquired in two separate batches, explaining the discrepancy in statistical significance for the ATF3^+^/CRX^+^ cells. **(d)** Receptor-ligand (R-L) analysis detailing ligands significantly upregulated in OSM-treated Müller glia whose corresponding receptors are expressed in at least 10% of rods or cones. Ligands are ranked by log_2_ fold-change relative to control. **(e)** Representative immunostainings (left) and quantification (right) of glial reactivity (p-STAT3, GFAP) and ER stress (ATF3) markers in mouse retinal organoids treated with LIF for 48h. Points represent independent organoid replicates; *\*\*\** p < 0.001, *\*\** p < 0.01; Cohen’s h = 0.7, 0.2, 0.4 for p-STAT3, GFAP, ATF3; scale bars: 100 *µ*m; insets, 20 *µ*m.OSM +

**Extended Data Figure 10.**
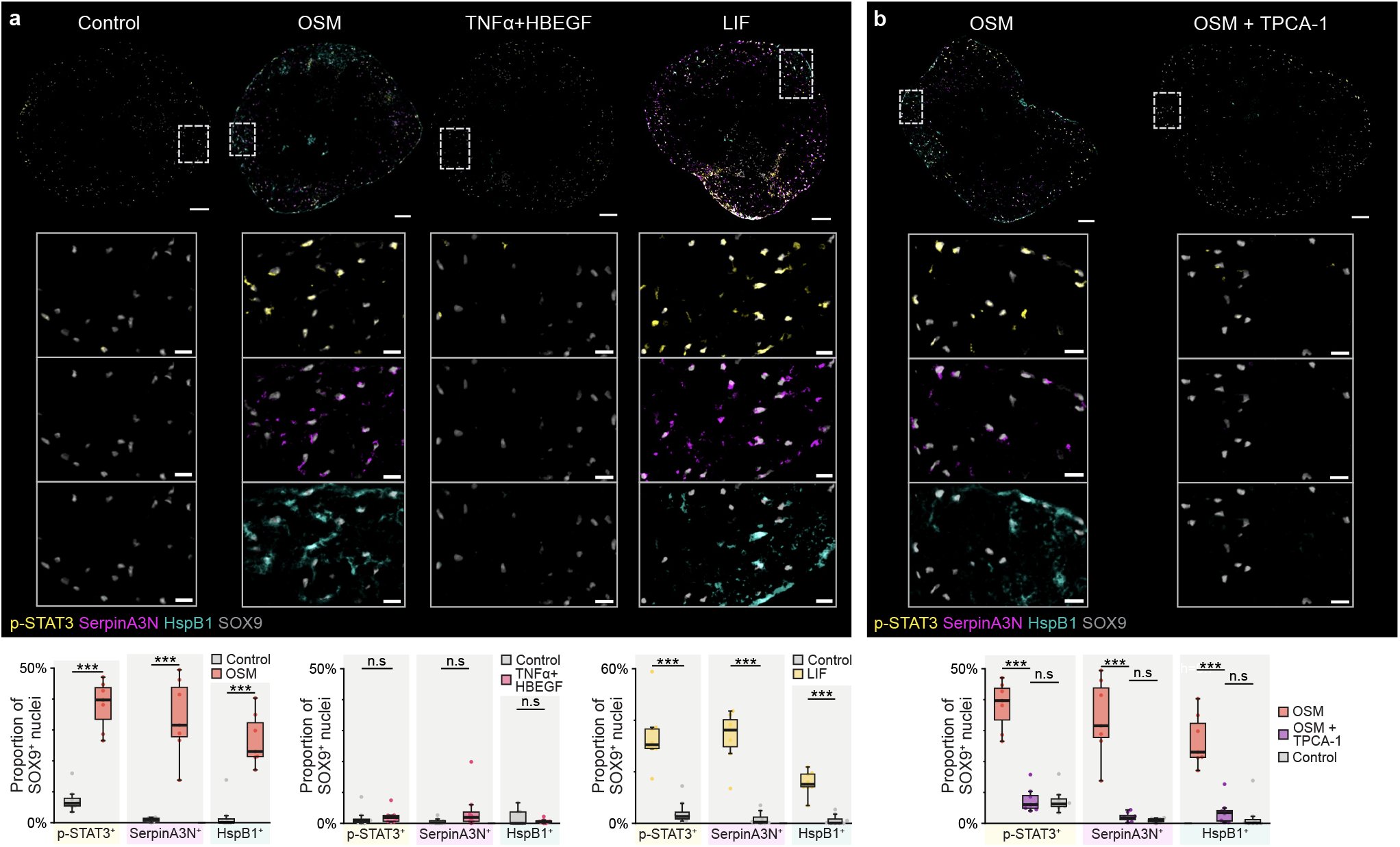
OSM but not TNF-*α* + HB-EGF induces brain astrocyte reactivity via the gp130–STAT3 signaling axis. **(a)** Representative immunostainings (top) and quantification (bottom) of glial reactivity markers p-STAT3, Serpin A3N, and HspB1 in mouse brain organoids treated with OSM (100 ng/mL; 48h), LIF (1000 U/mL; 48h), or TNF-*α*+ HBEGF (50 ng/mL; 120h). Respective controls are from identical time points and FFPE sections. Points represent independent organoid replicates; *\*\*\** p < 0.001, n.s for p > 0.05; Cohen’s *h*_p-STAT3_ = 0.8/0.8, *h*_SerpinA3N_ = 1.1/1.1, *h*_HspB1_ = 0.9/0.7 for OSM/LIF; scale bars: 100 *µ*m; insets, 20 *µ*m. **(b)** Representative immunostainings (top) and quantification (bottom) of astrocyte reactivity markers p-STAT3, Serpin A3N, and HspB1 in mouse brain organoids following co-treatment of OSM (100 ng/mL; 48h) with TPCA-1 (5 *µ*M; 48h). Points represent independent organoid replicates; *\*\*\** p < 0.001, n.s for p > 0.05; Cohen’s h = 0.8, 1.0, 0.7 for p-STAT3, SerpinA3N, HspB1; scale bars: 100 *µ*m; insets, 20 *µ*m.

## Supplementary information

**Supplementary Table 1**

Average presence scores per organoid dataset for each reference cell type or subtype, related to Fig. 1.

**Supplementary Table 2**

List of time points for each modality, related to Fig. 1, 2, and Extended Data Fig. 4, 5.

**Supplementary Table 3**

Selected doses by qualitative cytotoxicity assessment, related to Fig. 3, 4, 5, and Extended Data Fig. 6, 7, 8, 9, 10.

**Supplementary Table 4**

List of inflammatory cytokines used and their connection to CNS stress and disease, related to Fig. 3 and Extended Data Fig. 6.

**Supplementary Table 5**

In vivo gene signatures, related to Fig. 3, 4, 5.

**Supplementary Table 6**

List of targeted pathways and their relevance in retinal disease, related to Fig. 4, 5, and Extended Data Fig. 7, 8, 9, 10.

